# Learning immune cell differentiation

**DOI:** 10.1101/2019.12.21.885814

**Authors:** Alexandra Maslova, Ricardo N. Ramirez, Ke Ma, Hugo Schmutz, Chendi Wang, Curtis Fox, Bernard Ng, Christophe Benoist, Sara Mostafavi, the Immunological Genome Project

## Abstract

The mammalian genome contains several million cis-regulatory elements, whose differential activity marked by open chromatin determines organogenesis and differentiation. This activity is itself embedded in the DNA sequence, decoded by sequence-specific transcription factors. Leveraging a granular ATAC-seq atlas of chromatin activity across 81 immune cell-types we show that a convolutional neural network (“AI-TAC”) can learn to infer cell-type-specific chromatin activity solely from the DNA sequence. AI-TAC does so by rediscovering, with astonishing precision, binding motifs for known regulators, and some unknown ones, mapping them with high concordance to positions validated by ChIP-seq data. AI-TAC also uncovers combinatorial influences, establishing a hierarchy of transcription factors (TFs) and their interactions involved in immunocyte specification, with intriguingly different strategies between lineages. Mouse-trained AI-TAC can parse human DNA, revealing a strikingly similar ranking of influential TFs. Thus, Deep Learning can reveal the regulatory syntax that drives the full differentiative complexity of the immune system.

## INTRODUCTION

The immune system has a wide array of physiological functions, which range from surveillance of the homeostasis of body systems to defenses against a diversity of pathogens. Accordingly, it includes a wide array of cell-types, from large polynuclear neutrophils with innate ability to phagocytose bacteria, to antibody-producing B cells, to spore-like naïve T cells whose effector potential becomes manifest upon antigenic challenge. With the exception of rearranged receptors, all immunocytes share the same genome, and this phenotypic diversity must thus unfold from the genome blueprint, each cell-type having its own interpretation of the DNA code. This differential usage is driven by the interplay of constitutive and cell-type specific transcription factors (TFs) and regulatory RNA molecules, and possibly yet unknown sequence-parsing molecular entities.

Cell-specific recognition and effector potential are anchored in the cell’s transcriptome, itself a reflection of the conformation of DNA within chromatin which enables the expression of accessible genes, directly or as modulated by triggers from cell receptors and sensors. Recent technical advances reveal chromatin accessibility with high precision and across the entire genome ^1^, providing reliable charts of chromatin structure through immune cell-types ^2-5^. In these, Open Chromatin Regions (OCRs) reflected quite closely gene expression in the corresponding cells. The question, then, is to move from these descriptive charts to an understanding of how these chromatin patterns are determined. Analyzing the representation of TF-binding motifs (TFBS) in these differentially active OCRs provided some clues as to the TFs potentially responsible for cell-specificity, especially by using the cell-type specific expression of the TFs themselves as a correlative prior ^2^. Although motif analysis is a mature tool, it relies on imperfect TFBS tables assembled from different sources of data, with unavoidable noise, and more importantly does not provide insights into functional and cellular relevance of sequence patterns.

Artificial neural networks present a powerful approach that can learn complex and non-linear relationships between large sets of variables, and can recognize patterns whose combinations are predictive of multifaceted outcomes. Convolutional neural networks (CNNs) in particular can learn the combinatorial patterns embedded within input examples without the need for alignment of examples. Recent studies have begun to take advantage of CNNs to tackle aspects of gene regulation ^6^, including models that predict chromatin state ^7-9^, TF binding ^10,11^, polyadenylation ^12^, or gene expression ^7,13^ solely on the basis of DNA (100bp-1Mb) or RNA sequences, with the potential to ferret out relevant motifs.

The ImmGen consortium has recently applied ATAC-seq to generate an exhaustive chart (532,000 OCRs) of chromatin accessibility across the entire immune system of the mouse (81 primary cell-types and -states) ^2^. The data encompass the innate and adaptive immune systems, differentiation cascades of B and T lymphocyte lineages, detailed splits of myeloid subsets at baseline or after activation. We reasoned that it might provide the power to push the boundaries of what CNNs can learn, in terms of: (i) learning, solely from the DNA sequence of the OCRs, their pattern of activity across the immune system; (ii) extracting the sequence motifs, and their combination, that result in these predictions. The results showed that CNN model we derived (referred to as “AI-TAC”) can accurately predict the fine specificity of cell-type specific OCRs. The position of the uncovered motifs that are influential *in silico* recapitulated the binding sites of their molecular counterparts in “real” ChIP-seq data, and the motifs learned by AI-TAC are also a highly accurate match to known TF motifs, revealing novel aspects of the regulatory strategies used by different immune cells.

## RESULTS

### AI-TAC can predict enhancer activity from sequence alone

We developed and trained a deep convolutional neural network (CNN), hereafter AI-TAC, to predict the chromatin accessibility profiles across 81 immune cell-types on the basis of DNA sequences alone. In this way, AI-TAC learns the relationship between the combination of sequence motifs embedded within an OCR and its accessibility profile across immune cell-types. **Fig. 1a** illustrates the steps of training, interpretation and biochemical validation followed to ask whether a CNN model could effectively decipher, starting from DNA sequence, the motifs that drive the cell-specificity of immune gene expression.

**Figure 1.**
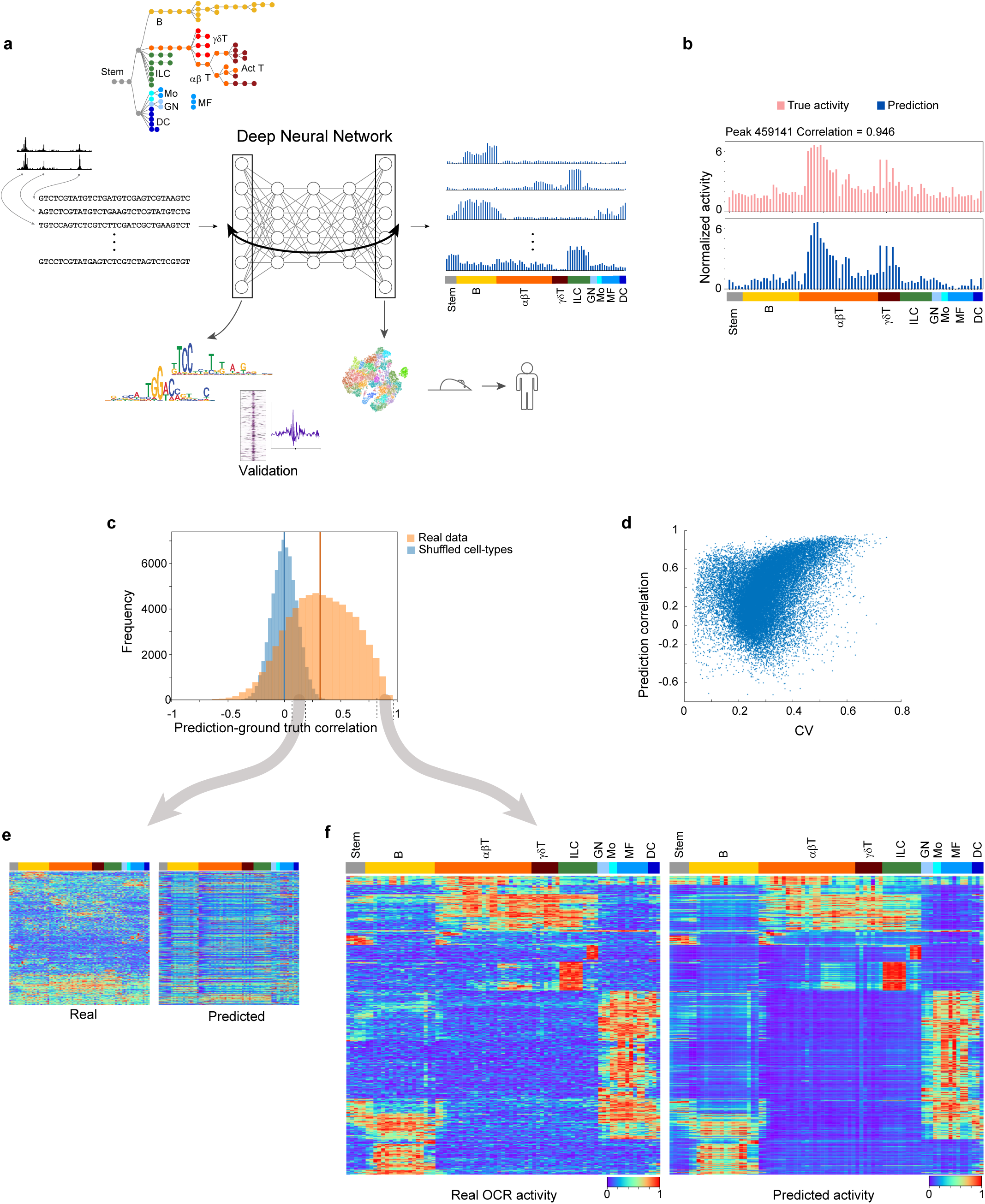
AI-TAC learns to predict cell-specific ATAC-Seq activity from sequence composition across the mouse immune system. **a**, A schematic of the AI-TAC model and its validation. AI-TAC is a deep convolutional neural network that takes as input OCR sequences and outputs ATAC-seq accessibility profile for 81 mouse immune cells. The sequence features (motifs) that are predictive of chromatin accessibility are learned during the training process. By analyzing the first and later layer filters, we derive important motifs and their combination that enable the model to make prediction for given OCRs. The predictions and motifs derived by AI-TAC are validated based on ChIP-seq datasets. **b**, Observed (top) and predicted (bottom) chromatin states of 81 immune cell-types for a single test OCR. **c**, Histogram of AI-TAC test set predictions trained on real data (orange) versus a model trained and tested on samples with randomly permuted chromatin accessibility profiles (blue). **d**, The coefficient of variation of the test set OCR chromatin accessibility profile on the x-axis versus the AI-TAC prediction correlation for those OCRs on the y-axis. **e-f**, Observed (left) and predicted (right) chromatin accessibility profile for real OCRs with **e** |corr| < 0.1 and with **f** corr > 0.8.

In practice, the model was trained by using as input 90% of 327,927 sequences, each 251bp long, of the OCRs defined by our recent ATAC-seq effort ^2^, to predict as output the profiles of ATAC-seq of each OCR across all measured cell-types using a multi-tasking strategy. The ability of the CNN to learn an accurate mapping between inputs and outputs depends on several hyperparameters, including the number of hidden layers, filters and their length. Bayesian optimization ^14^ showed that an architecture with three convolutional layers followed by two fully connected layers, with 19 bp sequence detected by the first layer filters, resulted in lowest achieved error on the validation data (**Fig. S1a-b**). We also found that the form of the loss function resulted in differential ability to predict cell-type specific activity profiles: using Pearson correlation as the loss function metric enhanced the ability of the model to accurately predict sequences whose activity varies across cell-types (p=10^−89^; **Fig. S1 c**,**d**). On a subset of held-back OCRs, the trained AI-TAC model showed impressive performance on precisely predicting granularly variable accessibility across all populations, as shown for one example **Fig. 1b**. Overall, 61% of test OCRs were predicted with a statistically significant correlation coefficient (at FDR 0.05, **Fig. 1c**). We observed largely monotonic relationship between the predictability of an OCR and its variation between immune cell-types, as OCRs with low prediction performance typically had small coefficients of variation (**Fig. 1d**). This relationship was expected given the mathematical implication of the loss function utilized, but this graph also indicates that the model is not missing out on particular classes of OCRs beyond that are ubiquitously active (as confirmed in the heatmap of **Fig. 1 f,g**).

We assessed the robustness of these predictions by performing several randomization experiments to create three different null models (**Fig. 1c, S2a**), as well as performing chromosome leave-out experiments with 19 different models (**Fig. S2b**). In addition, we performed 10 independent trials of 10-fold cross-validation (i.e., 100 trained models) so that each of the 327,927 OCRs was considered as part of ten different test sets (**Fig. S2c,d**). These data allowed us to confirm that well predicted OCRs were generally well predicted across different models trained on different subsets of the data, suggesting that regulatory logic captured by the model was generalizable.

### Learned motifs are associated with known pioneer factors and their lineage specificity

To understand the regulatory syntax learned by the AI-TAC model, we started by interpreting the first layer filters. Each filter in the first convolutional layer is “activated” by a short sequence, the sequence motif that activates each of these first layer filters being optimized during training. We first defined operational parameters of robustness for each of the 300 first-layer filters: *reproducibility* (how often its motif recurs in independently trained models), *influence* (how much it contributes to the prediction accuracy, measured by iterative nullification of each filter), and *frequency* (how many OCRs in the dataset activate it) (Methods, **Table S1**). We computed influence values on the set of well-predicted OCRs (with prediction correlation greater than 0.75), but confirmed that the results were similar when using only the test OCRs (**Fig. S2f**). Following the strategy of ^7^, we reconstructed the motif learned by each filter by finding the 19bp sequences (the filter length determined by Bayesian Optimization) within OCRs that are activated by each filter, determining their consensus (represented as a position weight matrix - PWM) and computing the *information content* (IC) in these motifs (**Table S1)**. For a baseline comparator, we applied the DeepLIFT framework ^15^ to these data and cross-matched AI-TAC results. DeepLIFT did not yield more determining motifs, once redundancy was accounted for, if anything less (Fig. S3).

Combining these parameters revealed two major groups among these trained first-layer filters (**Fig. 2a**): filters in the first group (e.g. 133, 167, etc.) were re-discovered repeatedly in every or almost every independent training run, had high influence (>10^−4^) and IC, with typically short (8-12 bp) consensus motifs reminiscent of typical TF binding sites. The second group (e.g. 259, 37, 249, 241) had far less reproducibility, influence and IC, with motifs that only included a few scattered bases or where less focused (∼15 bp long). Some of these low-influence and non-reproducible filters may represent noise in the neural network ^16^, or yet unknown regulatory motifs whose similarity structure may escape conventional alignment algorithms. We focused the rest of the analysis on the 99 reproducible filters, as a model retrained using these had only a small drop in performance as compared to the full model (**Fig S2e**, 99 altogether). As illustrated in **Fig. 2b**, reproducible filters partitioned between filters with restricted distribution (activating 10^3^ to 10^4^ OCRs) and generally higher influence, and a group of more frequently activated filters with overall lower influence and IC. To identify known motifs associated with the learned PWMs, we searched the Cis-BP database of TF motifs ^17^ using the TomTom algorithm ^18^. 101 of the 300 learned PWMs corresponded to at least one known TF motif at q-value < 0.05 (**Table S2**), and interestingly majority of these annotated PWMs belonged to the set of reproducible filters: 76 of 99 reproducible filters correspond closely to known TF motifs, many with astonishing similarity (as illustrated for Runx, Ets, and Ctcf in **Fig. 2c**). In 10 cases, the model also discovered exact reverse complements of the same motif (e.g. CTCF in **Fig. 2c)**.

**Figure 2.**
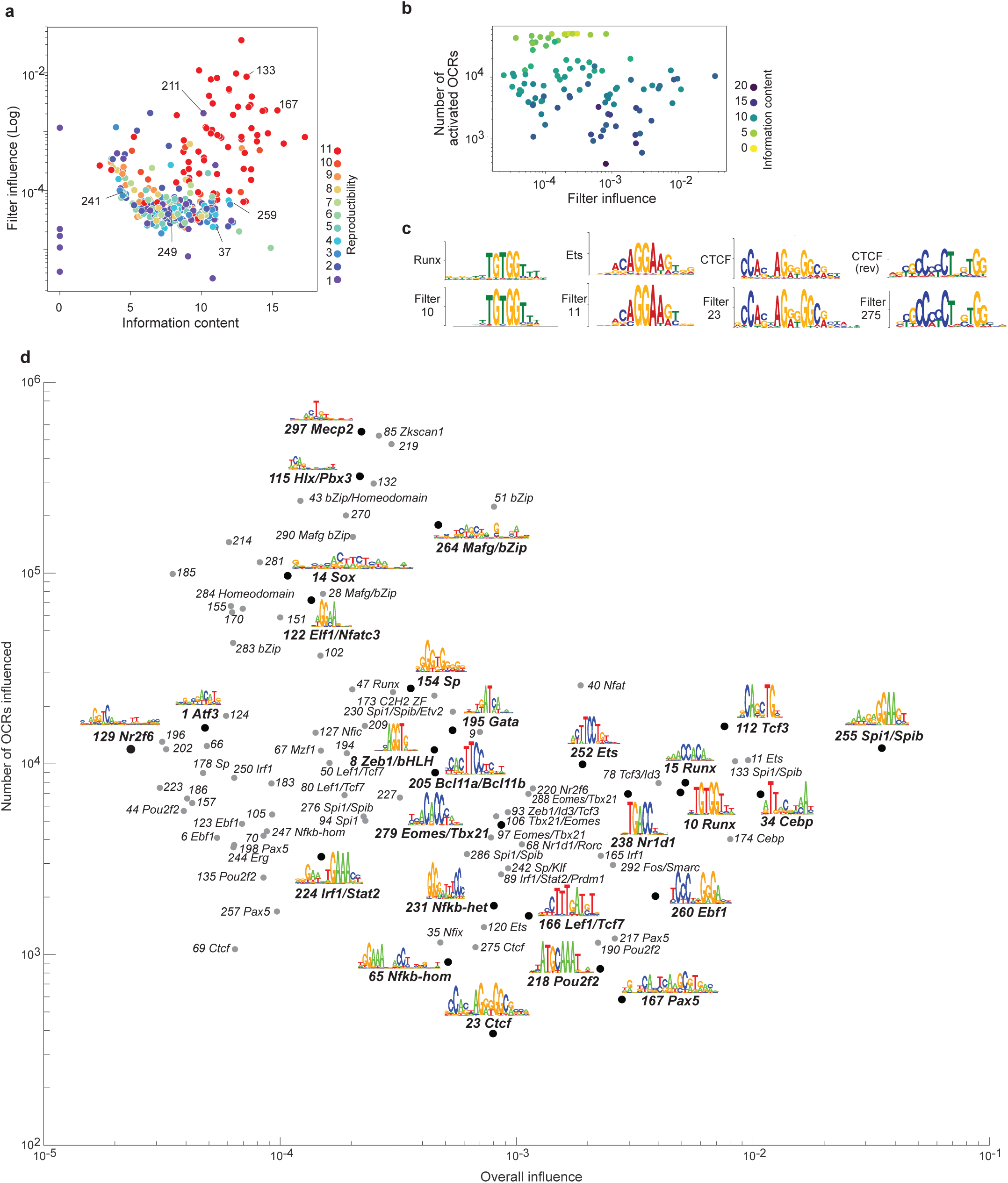
AI-TAC learns a wide range of motifs that together predict immune differentiations. **a**, The reproducibility of each of the 300 AI-TAC first layer filters across 10 additional re-trained models. Each model is trained on a different 90% subset of OCRs and initialized with different values. A filter is defined as “reproduced” in a different model if it matches any filter with a TomTom q-value of 0.05 or less. **b**, Log of filter influence versus log of number of OCRs activated by the filter and colored by information content of the filter’s PWM. An OCR is considered activated by the given filter if any of the filter activations for that OCR are at least ½ the maximum activation value of that filter across all input sequences, indicating the presence of that filter motif in the OCR. **c**, Examples of first layer filter PWMs and their alignment to known mouse TF motifs in the CIS-BP database, found using the TomTom alignment algorithm. **d**, Influence versus the number of OCRs with filter influence > 0.0025 for 99 filter motifs reproducible in at least 80% of model training iterations.

The regulatory landscape of chromatin opening throughout immune cell differentiation, as learned *de novo* by AI-TAC, can thus be summarized by the 99 motifs displayed in **Fig. 2d** (given the known complexities of TF motif assignments, that can reflect promiscuity and variation with cofactors or post-translational modifications, we opted for caution when several alternative TFs were candidates, annotating several filters at the family level only). We further refined the annotation of the most likely TF to each motif by combining Cis-BP scores with the correlation between activity of the OCR and expression of the TF across cell-types (illustrated for Pax5 in **Fig. S4a**); these correlations were comparable for filters annotated to the same TF (**Fig. S4b; Table S3**). The resulting set re-discovered several canonical regulators of lineage differentiation: Pax5, Ebf1, Spi1 (aka PU.1), Gata3. Other TFs were perhaps less expected in the context of cell-specific expression such as CTCF, a ubiquitous TF better known for its structural role in nuclear architecture ^19^, suggesting a cooperative role of CTCF as an adjunct to lineage-specific factors. A few influential TFs were represented by several filters, usually with slightly different motifs for the same TF (**Fig. S4c**). These nuances may correspond in part to technical noise from model over-parametrization ^16^, but they are also expected from known degeneracy in TF binding specificity, which is further influenced by interactions within dimers, as exemplified by the NF-κb family ^20^. AI-TAC filters indeed distinguished the canonical NF-κb hetero-dimer (filter231) and homodimer (filter247) motifs. For a broader perspective on how AI-TAC understands NF-κb family binding sites, we clustered the PWMs of all filters annotated to NF-κb through ten independent training runs. Interestingly, the hetero-dimer motif resurfaced regularly, while other motifs were less frequently discovered or allowed more sequence variation, suggesting gradations in their functional importance (**Fig. S4d**). Finally, several filters corresponded to motifs with no significant matches in Cis-BP or similar databases (**Fig. S5**). Some were short, and plausibly corresponded to half-sites, but others were longer and more complex, and may correspond to unrecognized TF binding motifs.

### Learned motifs associated with cell-type profiles

In addition to the overall influences, we next computed a per cell-type *influence profile*, quantifying the predictive importance of each filter in each of the 81 immune populations, as the difference between prediction values with and without each filter on a per cell-type basis (**Fig. 3a**). Interestingly, this analysis revealed both positive and negative influences (**Table S1**). Several of these positive influences (where the filter is needed for the full activation in the predicted profiles) were highly consistent with known roles of the corresponding TFs: Pax5 and Ebf1 essential for B cell differentiation, Spi1 and Cebp in myeloid cells, and Tbx21/Eomes in NK cells. AI-TAC was able to identify granular specificity of TFs beyond lineage-level importance: for instance, in the B lineage (top left of **Fig. 3a**), Pax5 showed pronounced influence early in proB stages, and late in germinal center B cells, while Pou2f2 (Oct2) was influential only in the latter. In myeloid cells, CEBP seems particularly influential in neutrophils, monocytes and tissue macrophages, but less so in dendritic cells (consistent with ^2^) and perhaps more surprisingly in central nervous system (CNS) microglia, an interesting notion given that microglia have a distinct origin from most other macrophage populations. No filter had the same degree of influence for T cells as Cebp/Spi1 or Pax5/Ebf1 had for myeloid and B cells.

**Figure 3.**
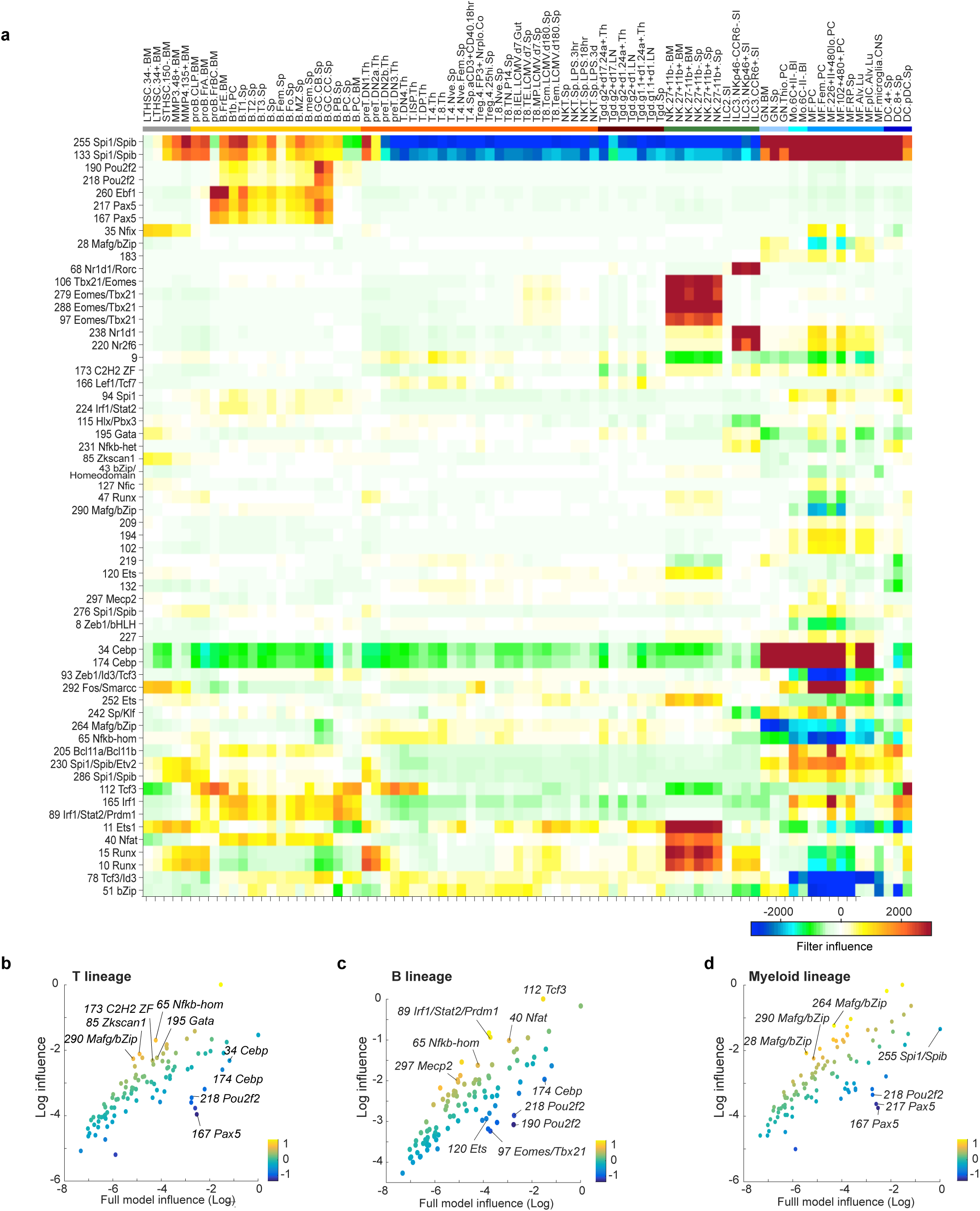
Cell-type profiles of learned motifs. **a**, Cell-type specific influence profile for the 99 reproducible filters found in at least 80% of model training iterations. **b-d**, Log of filter influence in original AI-TAC model versus AI-TAC fine-tuned for one additional epoch exclusively on the **b**, T-cell lineage, **c**, B-cell lineage and **d**, myeloid cell lineage.

More paradoxical were the *negative* influences (where the predicted activity of OCRs is over-estimated in its absence). These occurred most prominently for the myeloid-specifying motifs recognized by Spi1 or Cebp, but not for every strong filter (i.e. not for Pax5 or Tbx21 motifs). Thus, the neural network uses the presence of a Spi1 motif in an OCR to enforce its inactivity in T cells, beyond the neutrality that might be expected from a missing TF, a concept possibly connected to the need for Spi1 shut-down for T cell differentiation ^21^.

AI-TAC appears to learn stage-specific influences within lineages. To go deeper in identifying motifs and TFs that have predictive influence within a lineage, we used a “fine-tuning” strategy ^22^: the AI-TAC model (trained for 10 epochs initially) was further retrained for one epoch to predict chromatin accessibility in only the T, or B or myeloid datasets (**Table S4**). Such a fine-tuned model should reconfigure its learned parameters to enhance its ability to predict the cell-type-specific accessibility profile within a given lineage. When comparing the filters in these fine-tuned models to the initial AI-TAC model, we observed that the CNN selectively down-weighted (“forgot”) filters in a very logical manner, e.g. decreased influence of B-specific Pax5/Pou2f/Ebf1 filters in T or myeloid models, or of Cepb filters in non-myeloid models (**Fig. 3b-d**). Conversely, several filters gained influence in all three (Tcf3 (aka E2A), NFkB, Zkscan1, Maf, bZIP). This is consistent with the notion that E2A or NF-κb are more involved in modulating differentiation or activation states within all immune lineages, rather than specifying them. Because the dataset included 22 distinct T lineage cells, providing sufficient training data, we also trained an AI-TAC^T^ model solely on T cells (**Fig. S6a**). Many of the same filters reappeared, but with a slightly different emphasis (e.g. the influence of NF-κb is high during pre/pro-T stages but wanes in resting mature T cells, but returns in Tregs and activated/memory CD8+ T cells).

If AI-TAC accurately predicts cell-specificity of OCRs activity, it should also correctly assign differences in chromatin accessibility between cells. This was clearly the case for the OCRs that differ between T cells and macrophages (**Fig. S6b**). We also tested predictions for OCRs whose activity is induced by innate receptor triggering (NKT cells, 3 hrs after LPS injection *in vivo*, **Fig. S6c**). There was again a very good fit between predicted and observed OCR activity, in particular for OCRs influenced by NF-κb filter231 (NF-κb-het) and whose accessibility is upregulated by LPS, *in vivo* and *in silico* (**Fig. S6c**). Thus, AI-TAC can faithfully map differences between cell-types, for large inter-lineage distances as well as more focused responses to cell activation.

### Biochemical validation of predicted TF binding

While the identity of the motifs learned *in silico* were striking, and the distribution of their influence across cell-types made sense, it was important to validate the significance of these observations. We first selected the 500 OCRs most influenced by filter167 (Pax5), which aligned on the Pax5 consensus motif (**Fig. 4a**). In accordance with expectations, these OCRs were active in B cells, but not in thymic DPs (**Fig. 4a**, right panels). We then examined the fit between the in silico learned filters and the actual position of the corresponding TFs in the genome, deduced from chromatin immuno-precipitation (ChIP-seq). Overall, we observed a very strong concordance between AI-TAC’s predictions and ChIP-seq data. As one example, OCRs predicted to be influenced by filter 255 (Spi1) recapitulated the two main binding sites of Spi1 in the *Il1b* locus (**Fig. 4b**). More generally, the top OCRs influenced by filters 167 (Pax5), 260 (Ebf1) or 166 (Lef1/Tcf7) strikingly overlapped with binding sites defined by ChIP-seq for those factors, relative to control OCRs (0.006 to 0.09) (p<0.003, **Fig. 4c**).

**Figure 4.**
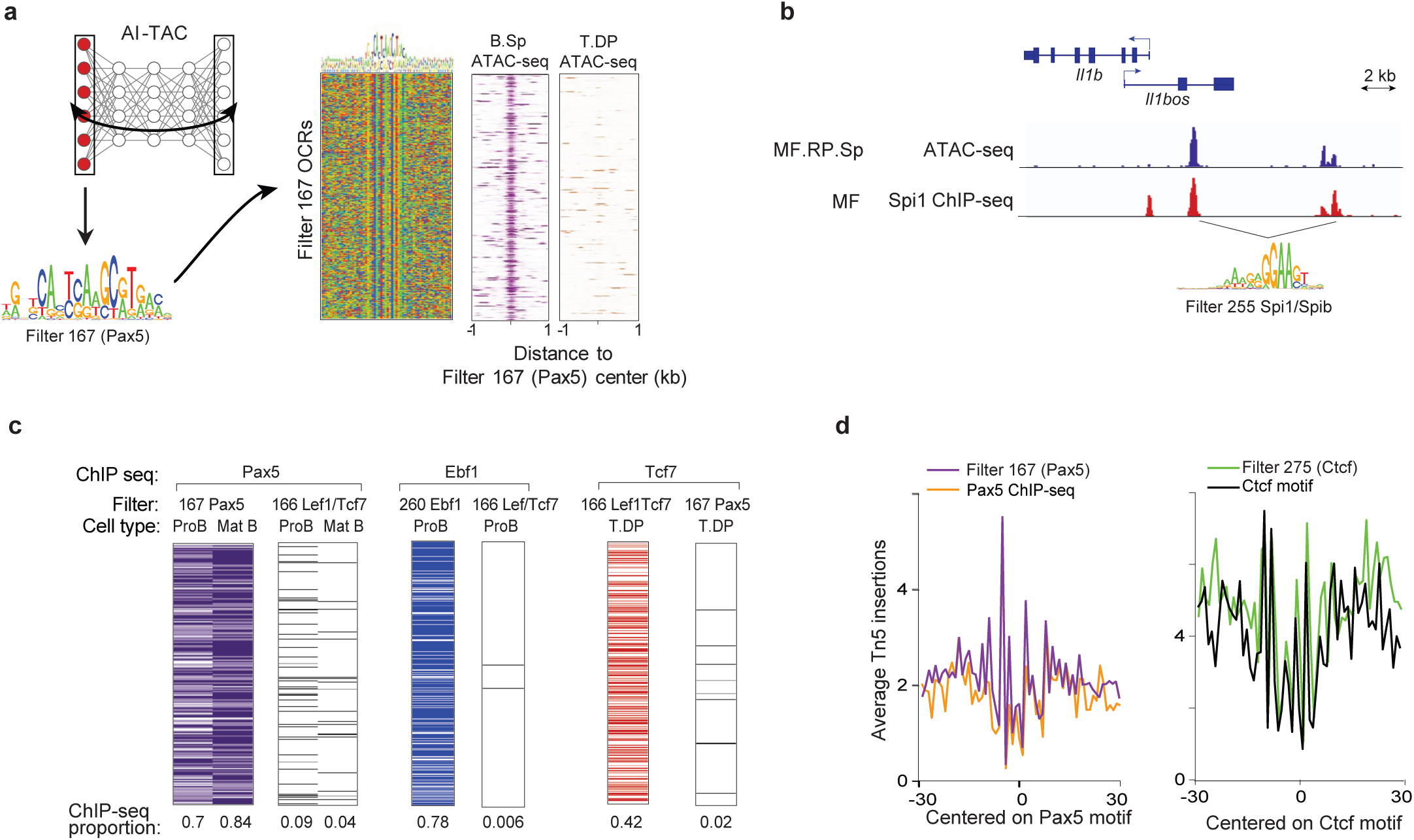
Biochemical validation of AI-TAC learned motifs. **a**, The top 500 OCRs influenced by filter167 (Pax5) were selected, their consensus verified (middle panel) and their ATAC signal in B cells or thymic DPs displayed at right. **b**, Filter 255 activations indicate the presence of a Spi1 motif within two OCRs with known Spi1 binding sites in the Il1b locus. **c**, Validation of predicted Pax5, Ebf1, and Lef1/Tcf7 filters with *in vivo* ChIP-seq. Proportion of TF peak overlap was computed from influential OCRs (top 500). TF filter specificity was controlled by cross-comparisons of binding enrichments with AI-TAC predicted filters. Left to right; Pax5 bound in filter 167 (Pax5) OCRs pro (0.7) and mature B cells (0.84), cross-compared with filter 166 (Lef1/Tcf7) OCRs pro (0.09) and mature B (0.04). Ebf1 bound in filter 260 (Ebf1) OCRs Pro B (0.78) and filter 166 (Lef1/Tcf7) OCRs Pro B (0.006). Tcf7 bound in filter 166 (Lef1) OCRs T.DP (0.42) and filter 167 (Pax5) OCRs T.DP (0.02). **d**, AI-TAC predicted ATAC-seq footprint from Pax5 (167) OCRs, independently derived Pax5 ChIP-seq footprint, predicted footprint of CTCF (275) and CTCF motif.

Finally, we analyzed deep ATAC-seq traces in B lymphocytes at nucleotide-level resolution, where one can discern a “footprint” where the binding of a TF prevents or favors accessibility by the Tn5 transposase ^23^. We superimposed deep (>200M reads) ATAC-seq traces at positions predicted to activate Pax5 or CTCF filters in AI-TAC over true binding sites independently determined known from ChIP-seq and motif identification. Here again, AI-TAC driven predictions accurately coincided with true TF binding, showing the same fine details of accessibility (**Fig. 4d**). Thus, whether in matching the distribution of TF binding or the nucleotide-level traces to biochemically determined ones, AI-TAC *in silico* predictions are strongly validated by *in vivo* data.

### Dissecting the combinatorial logic of chromatin opening

That enhancer elements tend to occur as repeats has long been a theme, either because those first discovered in viral genomes occurred as tandem repeats, or because synthetically engineered enhancers were more effective as strings of the same motif. It was thus of interest to ask whether repeats of the same motif were enriched among active OCRs. This was not the case (**Fig. 5a**): there was no greater frequency of recurrence of activating filters within an OCR than would be expected by chance, with two interesting exceptions: filter231 (NF-κb-het), consistent with the demonstration that NF-κb uses clustered binding sites non-cooperatively to incrementally tune transcription ^24^; and two GC-rich motifs recognized by Sp (242, 154), consistent with reports that SP1 functions best in the context of repeated GC-rich blocs ^25^. Interestingly, the relative positioning of these repeats was not random, e.g. maxima at 90 and 140 bp (nucleosome length?) for NF-κb (**Fig. 5b**). On the other hand, activation of the same filter by OCRs likely to control the same gene, as evidenced by regression ^2^ showed a significant enrichment compared to chance (**Fig. 5c**). Thus, shortly spaced repeats of controlling motifs are not a regulatory strategy commonly employed to control mammalian immune cell differentiation, motif repetition being provided by spaced elements connected by DNA loops.

**Figure 5.**
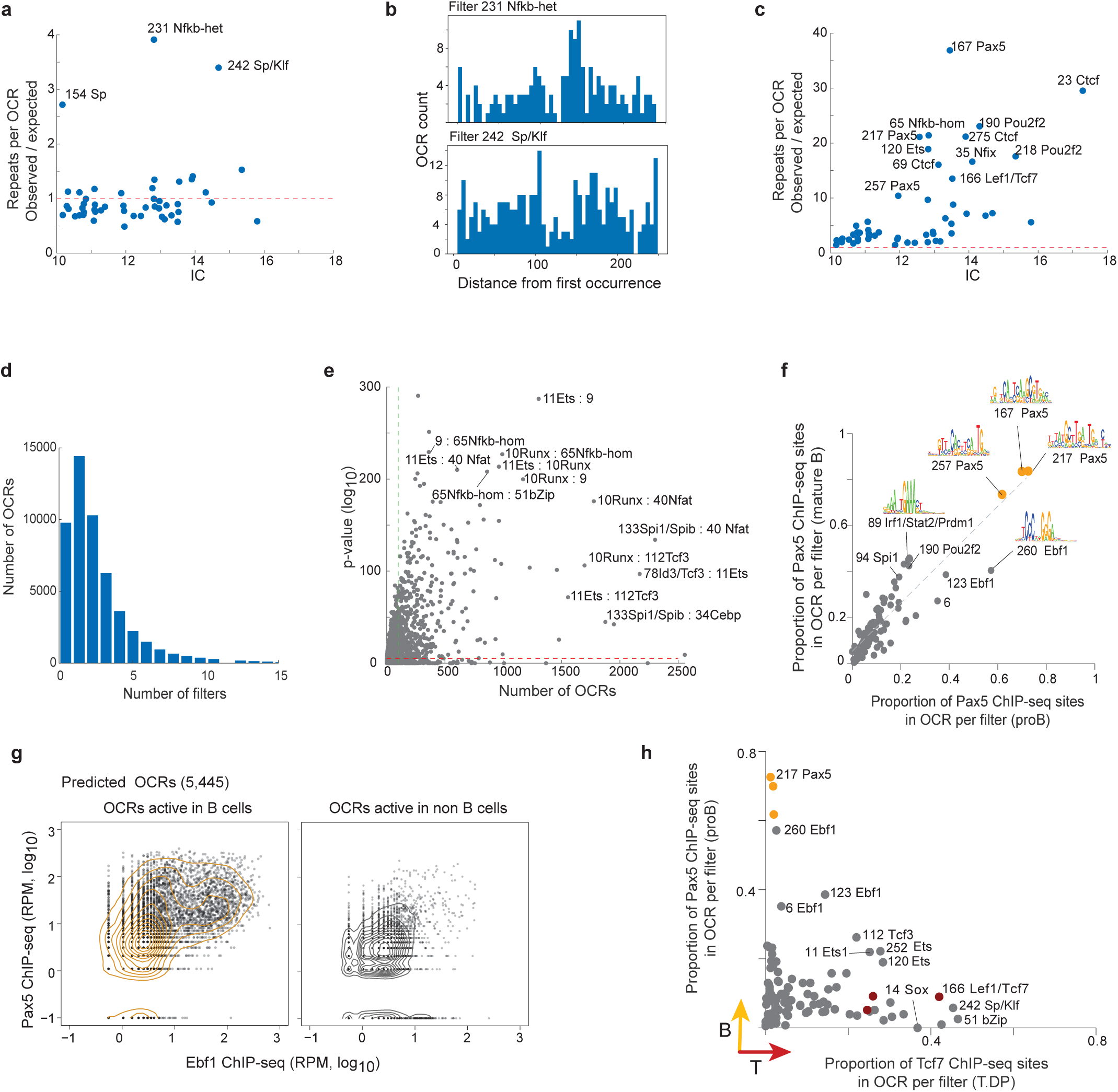
Identifying combinations of motifs that are predictive of immune differentiations. **a**, Each dot represents a filter. Y-axis shows the expected vs observed ratio for the number of OCRs that contained more than one instance of a filter’s motif. An OCR is defined as containing a filter motif if any of the activations for that OCR (across its 251bp length) are above ½ the maximum activation of that filter across all input sequences. **b**, Histograms show the distribution of distances between two occurrences of the same motif on a given OCR. **c**, For each filter, scatter plot shows the expected versus observed ratio for the number of genes whose set of assigned OCRs contained multiple instances of the same filter motif (y-axis) against the filter motif’s information content (x-axis). **d**, Histogram shows the number of filters per OCR that have an influence value of 0.0025 or more, which corresponds to a 5% impact on the correlation of the prediction. **e**, For each pair of filters, the number of OCRs where both filters were deemed influential is shown on the x-axis, and the hypergeometric p-value for the significance of the number of shared OCRs, as compared to expectation based on prevalence alone, is shown on the y-axis. To eliminate technical artifacts, filter pairs whose motifs were similar to each other (PWMEnrich>0.5) were removed. **f**, Enrichment of OCRs (top 500 influential OCRs per filter, n=49,500 OCRs) bound by Pax5 in AI-TAC reproducible filters (n=99) for pro and mature B cells. **g**, *In vivo* ChIP-seq occupancy for Pax5 and Ebf1 in AI-TAC predicted B cell OCRs (n=5,443). Co-occupancy patterns observed in predicted B OCRs and for non-B predicted OCRs (n=5,443). **h**, Enrichment of OCRs (top 500 influential OCRs per filter, n=49,500 OCRs) bound by Tcf7 or Pax5 ChIP-seq in AI-TAC reproducible filters (n=99) for T.DP and pro B cells respectively.

Given the size of the vertebrate genomes, combinations of transcriptional regulators are the only practicable solution to encode the complexity of development and cell-type differentiation ^26^. Pervasive interactions between TFs within multimolecular complexes have been observed in genomic and functional experiments, but an overall perspective on the combinatorial interactions that actually influence transcription remains incomplete. It was thus of interest here to ask which combinations of motifs are co-influential in AI-TAC’s predictions, and hence suggest co-operation between TFs. Because the higher order relationships between first layer motifs are encoded in the deeper layers of the network, an obvious first attempt at identifying important filter combinations is to look for combinations of motifs assembled by the second layer convolutional filters ^12^. Due to the maxpooling applied to the first layer output, constructing clear motifs from second layer activations was not possible and we instead examined the second layer filter weights directly. We found that in a large number of cases the second layer filters recognized similar (or reverse complement) first layer motifs, indicating that the second layer is perhaps assembling cleaner versions of first layer motifs rather than learning combinatorial logic (**Fig S7**).

As an alternative, we identified for each OCR the set of filters that impact the accuracy of its prediction (i.e., influence) by 5% or more. Of the set of OCRs that were influenced by at least one filter at this threshold, many (N=23,910, 56%) were influenced by 2 to 6 filters, and a few (N=1,514, 4%) were even impacted by 10 or more filters (**Fig 5d**). This large set of OCRs impacted by multiple filters provided a rich base to identify common co-influential motifs. To identify influential combinations between different TFs, we computed for each filter pair the number of OCRs that they both impact, and compared it to expected co-influence based on each filter’s prevalence. This analysis yielded 493 co-influential filter pairs (adjusted p<0.05, and number of co-occurrences>100) (**Fig. 5e, Table S5**). Interestingly, filters that are broadly influential tended to be significantly co-influential with each other (e.g., Ebf1 and Pax5, n=193, p<10e-20; Lef1/Tcf7 and Runx, n=471, p<10e-50). Among over-represented pairs, some TFs were highly recurrent, acting as “hubs” of sorts: Tcf3 (filters 78/8/93), Runx (filter10), Ets (filter11), and Nfat (filter40) co-occurred with 40 or more other filters (**Table S5**).

Some of these inferences in terms of motif co-influence were congruent with existing knowledge (e.g. Tbx21/Runx, Spi1/Cebp, etc.), but to provide proof-of-principle validation we again turned to ChIP-seq data. Using Pax5 ChIP-seq datasets generated in pro B and mature B cells ^27^, we asked what fraction of the OCRs influenced by each AI-TAC filter overlapped with a validated Pax5 binding site. As expected from **Fig. 4c**, OCRs influenced by filters 167, 217 and 257 (all annotated as Pax5) contained a high proportion of true Pax5 binding sites in both proB and mature B cells (0.62 to 0.83, **Fig. 5f**). Interestingly, OCRs influenced by several other filters also contained a high proportion of Pax5-binding sites (in particular filters 260 (Ebf1), 89 (Irf1/Stat2/Prdm1) or 190 (Pou2f1). Finding Ebf1 associated with Pax5-annotated filters is consistent with the known molecular collaboration between Ebf1 and Pax5 in controlling B-cell identity ^28-30^. This conclusion was borne out by displaying Pax5 and Ebf1 ChIP-seq signals in OCRs active in B cells, showing that some OCRs preferentially bound one of these two TFs, and many both (**Fig. 5g**).

As another validation and to further identify combinatorial signals, we compared ChIP-seq signals for Pax5 and Tcf7 (active in T cells) in OCRs predicted to be activated by different filters. OCRs influenced by Lef1/Tcf7 filters (166, 50, 80) were again strongly enriched in Tcf7 bound sites in DPs ^31^ but had low Pax5 signals in proB cells, while OCRs that activate Pax5 filters and its associated Ebf1 and Pou2f filters were enriched in Pax5 ChIP-seq signals in proB cells but low in Tcf7 (**Fig 5h**). OCRs that activated filters annotated to Sp1 (242) or bZip (51) were enriched in Tcf7 ChIP-seq, confirming that these TFs interact with Tcf7 (**Table S5**). Interestingly, AI-TAC predictions recovered regulators Tcf3/E2a (112) and Ets (11, 252, 120) with similar enrichments in Pax5 and Tcf7 bound sites, consistent with known overlapping regulatory function in the specification and maintenance of B and T lineages ^32^. Thus, combining AI-TAC predictions with *in vivo* ChIP-seq data parsed TF binding patterns with regulatory co-influence at different stages of T or B differentiation, and resolved novel regulatory motifs represented in Tcf7 bound sites across disparate T cell states.

### TF cis-regulatory syntax embedded in AI-TAC’s fully-connected layer

The last fully-connected layer of a neural network represents the final non-linear embedding of the input examples in the derived feature space. To visualize this space, we represented each well-predicted OCR by its activation values across the 1000 neurons of the last layer, and projected these activation vectors in 2-D using the t-SNE algorithm (**Fig. 6a**). When OCRs in this space were colored by their accessibility in different lineages, lineage-specific activity mapped to different segments (**Fig 6b**), indicating that this last layer discriminates well between lineages. Next, we analyzed how the influence of individual first-layer filters (and corresponding TFs) projected in this space. The influence of Pax5 (filter167) and Ebf1 (filter260) was highest in closely related poles of the B cell area, overlapping partially (**Fig. 6c**), in accordance with **Fig. 5f**. Similarly, the influence of Spi1 (255) and Cebp (34) in myeloid lineage OCRs was distinguishable, with some OCRs influenced by both (**Fig. 6d**), consistent with the known cooperativity of Spi1 and Cebp across myeloid cell-types ^33^.

**Figure 6.**
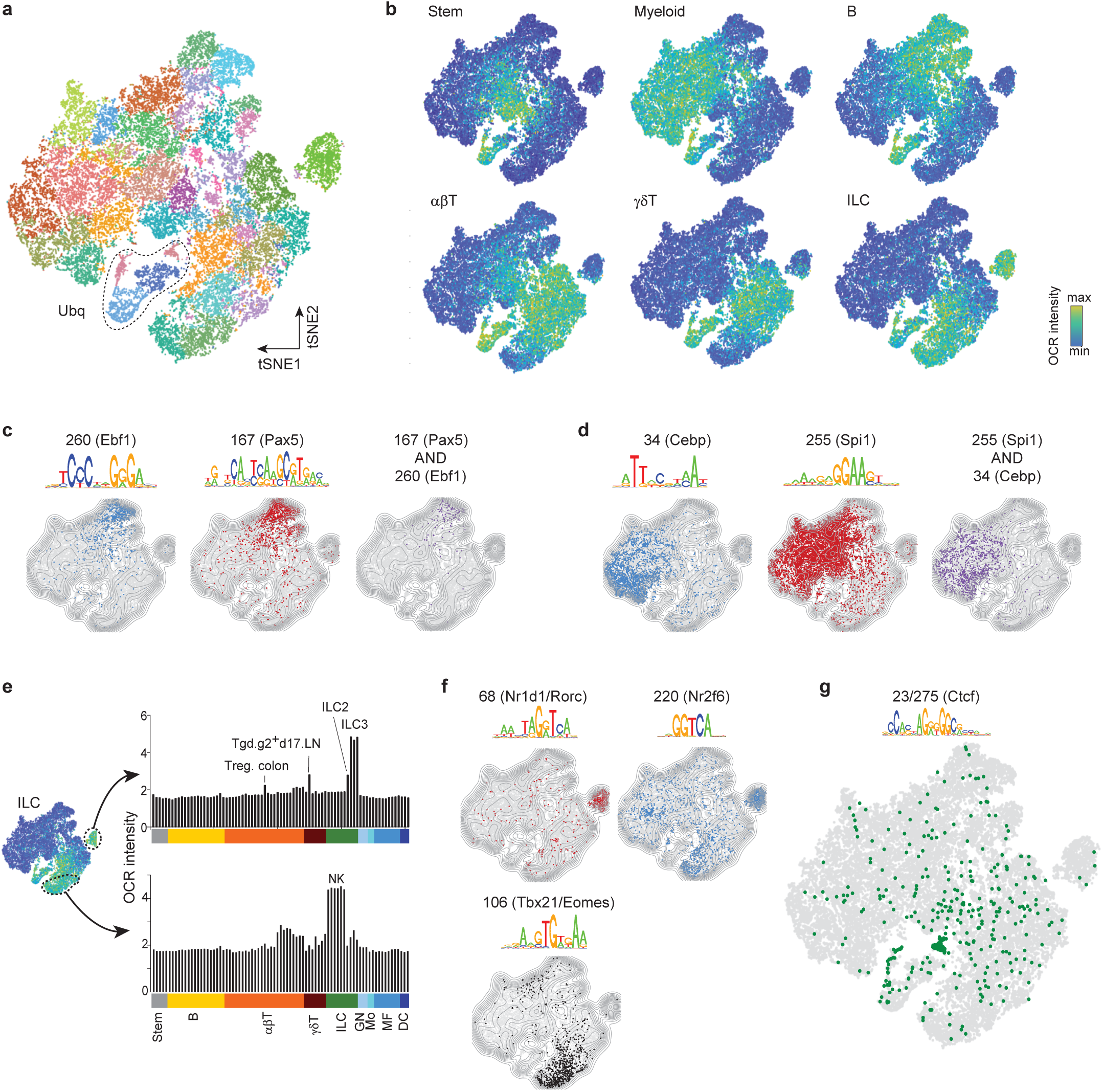
Identifying combinatorial regulatory syntax embedded in AI-TAC’s fully-connected layer. **a**, t-SNE representation and clustering of well-predicted OCRs (n=30,875) based on their scores across the last layer (695 nodes) of the trained AI-TAC model. **b**, ATAC-seq intensity of OCRs across immune lineages. **c**, OCRs influenced by filters 167 (Pax5), 260 (Ebf1) and co-occurring. **d**, OCRs influenced by filters 34 (Cebp), 255 (Spi1) and co-occurring. **e**, ILC lineage OCRs parsed by ATAC-seq mean intensities of ILC2/3 and NK subsets. **f**, OCRs influenced by filters 68 (Nr1d1/Rorγ, 220 (Nr2f6), and 106 (Tbx21/Eomes). **g**, OCRs influenced by filter 23 and 275 (Ctcf).

Among the patterns of OCR activity projected in this embedding space, the stratification of OCRs accessible in ILCs was intriguing (**Fig. 6b**), as it demarcated a cluster of OCRs distinct from all others. We cannot formally rule out that this demarcation of ILC3-active OCRs results from a technical artefact, although have no indication in this sense, but the dichotomy turned out to reflect a partition between OCRs active in NK cells vs ILC3 (and to a lesser extent in ILC2, colonic Treg and some Tgd cells; **Fig. 6e**). OCRs active in NK cells were influenced by Tbx21/Eomes-related filter106 (**Fig. 6f**), but ILC3-preferential OCRs were mainly influenced by filters annotated to the Nuclear Receptor (NR) family Nr1d1/Rorγ (68) and Nr2f6 (220) (**Fig. 6f**, see also **3a**). The influence of these NRs is consistent with the demonstration of a role for Nr1d1 in Ilc3 differentiation ^34^. Thus, chromatin activity and TF control learned *in silico* appear very different for these groups of ILCs.

Apart from predicting lineage specific patterns, the last layer also parsed a subset of OCRs with widespread activity across all lineages (“Ubq” in **Fig 6a**). These small clusters were characterized by the influence of the ubiquitous TFs Sp/Klf (filter242) and CTCF (filter23/275, **Fig 6g**), suggesting common structural motifs. Interestingly, the influence by CTCF filters was also observed in clusters of more cell-type specific OCRs, a disposition consistent with the notion that CTCF partakes in the generic organization of DNA topologies in the nucleus, but also cooperates with cell-type specific TFs to form specific loop and domain structures ^19^. Thus, AI-TAC’s final layer embedding of OCRs had the ability to refine lineage and cell specificity through TF influence patterns, suggesting that marginal influence estimates can serve as a proxy for the biological regulatory impact.

### Gene regulatory network inference based on predicted TF binding sites

We next assessed the relationship between chromatin accessibility, cis regulatory syntax at genes’ OCRs, and gene expression. Using results of a previous analysis in which we assigned *cis* OCRs to genes by correlating gene expression and chromatin accessibility across the 81 cell populations ^2^, we extrapolated from TFs that are influential for a given OCR in the AI-TAC model to sets of TFs predicted to be influential for regulation of each gene, generating a network of regulator TF to gene edges for 4,569 expressed genes (**Table S6, also available from www.immgen.org**). The median number of regulators per gene was three, but 32% of genes were assigned 5 or more regulators (**Fig. S8**). Fittingly, we observed a significant overlap (N=144, hyper-geometric p<10^−20^) between edges inferred in this analysis and those previously inferred from co-expression within independently generated microarray data ^35^.

The main TF hubs in the co-expression based regulatory network also dominated the rankings based on influence in the AI-TAC. For instance, Spi1, Ets, Tcf3 emerged as regulators of a large number of genes (**Fig. 7a**). To go beyond general lineage differences, we analyzed cell-type specific networks in the B, T, and myeloid lineages, by identifying differentially expressed genes and exploring their regulators. Clustering of genes based on their expression carried over only weakly to their assigned regulators (**Fig. 7b**), with considerable heterogeneity in the inferred networks and scattered participation of many other possible regulators. There were also differences in network structure between lineages: B and myeloid networks consisted of several interactive hubs (Ebf1, Pax5, and Pou2f2 in B; Spi1 and Cebp in myeloid), but the T cell network was more distributed, with major hubs not as apparent, consistent with **Fig. 3a**. In summary, this representation departs from the “modular structure” which considers gene expression reducible to a set of co-regulated genes. Instead, though each lineage is associated with major regulators, these regulators interact with a diverse set of TFs depending on the actual target gene.

**Figure 7.**
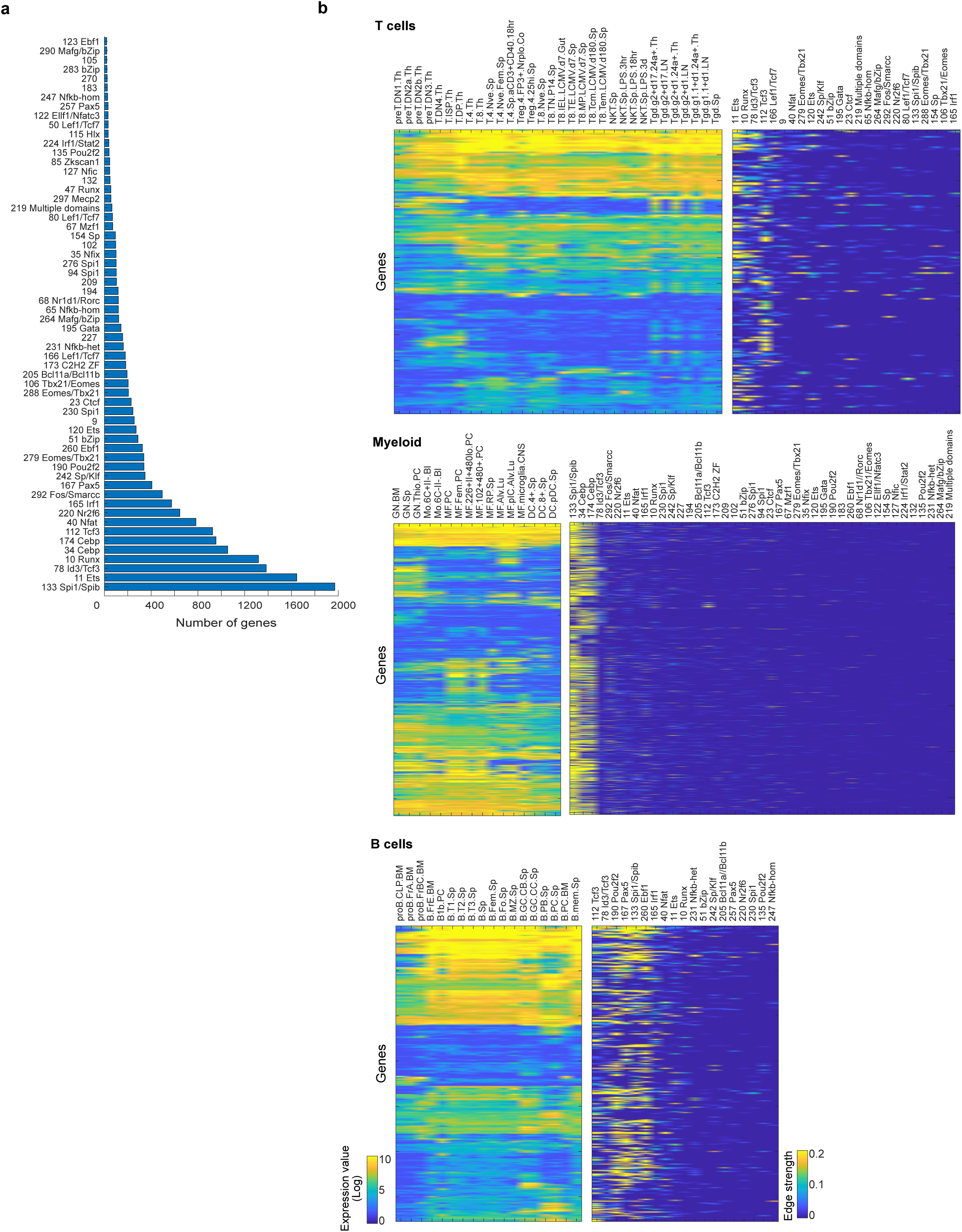
Combinations of RNA-seq expression data and OCR-motif predictions enables construction of gene regulatory networks. **a**, Histogram of the number of gene targets for each TF. **b**, For each lineage, RNA-seq based gene expression values for differentially expressed gene across cell populations is shown on the right and inference of TF assigned to each target gene is shown on the left. For each lineage, the ordering of genes (rows) is preserved between the two heatmaps.

### Cross-species generalization of AI-TAC predictions

The ultimate test of generalizability of a trained machine learning model requires assessing its performance on independent/external datasets. Because the human and mouse immune systems share many regulatory nodes ^36-38^, and TFs and their motifs are conserved across far wider evolutionary distance, we used cross-species testing to assess AI-TAC predictions on unseen human OCRs defined by a prior ATAC-seq analysis in 25 hematopoietic cell-types ^5^. We first used the ImmGen pipeline to pre-process the human dataset, identifying 539,611 OCRs of 251bp length. We then directly applied the mouse-trained AI-TAC model on these human sequences, and predicted their accessibility across the 8 cell-types from the mouse model that had a counterpart in the human dataset (**Table S7**). The correlation between predicted and observed accessibility profiles was significant for the majority of these OCRs, and highly skewed as compared to shuffled human OCR sequences (**Fig. 8a**).

**Figure 8.**
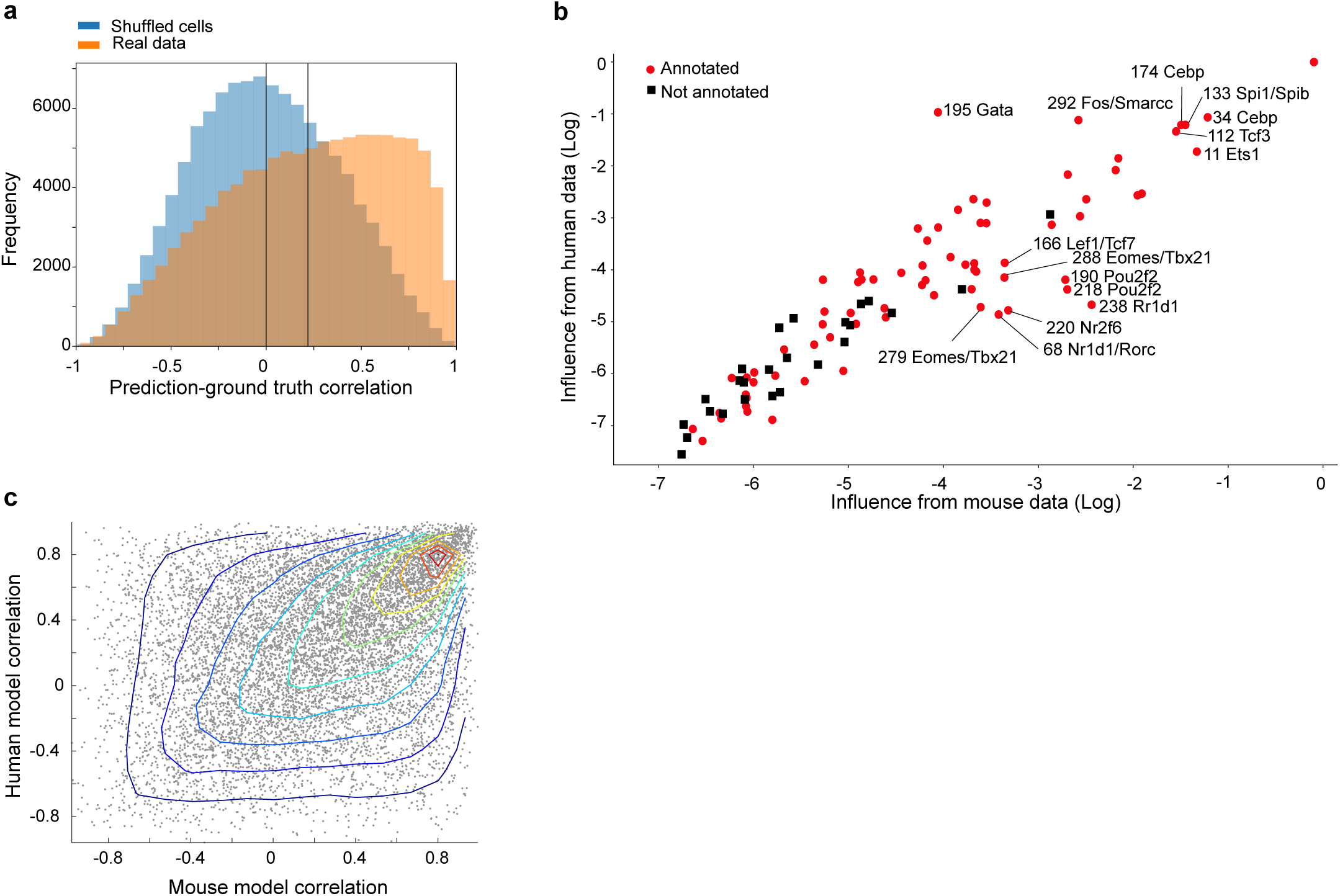
AI-TAC model is predictive of human OCR accessibility profiles. **a**, The trained AI-TAC model was directly applied to predict accessibility profile of human sequences underlying OCRs across eight cell-types that overlapped between mouse and human datasets. Figure shows histogram of AI-TAC predictions (measured by Pearson correlation between observed and the model’s predictions) on “real” human 251bp sequences underlying 539,611 OCRs (orange) versus randomly permuted human OCR sequences. **b**, Log influence of AI-TAC’s filters in mouse (x-axis) and human (y-axis), on the basis of nullification of each filter at a time. **c**, Prediction performance (measured by Pearson correlation) for each human OCR based on AI-TAC trained on mouse data (x-axis) and a model directly trained on human ATAC-seq data (y-axis). The human-based model was trained on 50% of human OCRs. The scatter plot only shows the performance of 50% of human OCRs that were considered part of the test set.

We then explored the degree of conservation of the important AI-TAC TF motifs. After fine-tuning the AI-TAC model on human data for four epochs decided by an early stopping strategy we obtained influence scores for each filter based on its prediction performance on the set of well-predicted human OCRs. We observed a striking correlation in terms of predictive influence of a filter in mouse and human datasets, indicating preservation of overall regulatory impact (**Fig. 8b**). Only a few outliers were noted, for example Gata, which in this case may be explained by the addition of erythroid cells in the human dataset.

Finally, to assess whether there exist broad classes of human OCRs that are not predictable by the mouse model, we trained a CNN directly on the human dataset and compared its prediction performance over the test set of OCRs to the mouse AI-TAC model. We observed a strong correlation between prediction performance of the mouse and human models on these human OCRs (**Fig. 8c**), with only a minor shoulder of OCRs that were better predicted by the human-trained model. This indicates that the regulatory code that is predictive of immune cell chromatin activity is extraordinarily conserved between human and mouse species.

## DISCUSSION

Differentiated cell states and functions, deeply encoded in the DNA sequence, unfold through the coordinated action of TFs and of the transcriptional modifiers they coopt. We show here that an artificial neural network can emulate these biological decoders and predict, based on sequence alone, cell-specific patterns of chromatin accessibility across the entire immune system. It does so with astonishing precision for many OCRs, fully matching biochemical validation data, and in the process re-discovering with extreme fidelity the binding sites for known transcription factors. By probing the sequence cues that the CNN detects and integrates, we infer how the biological decoders may operate, yielding a broad portrait of the sequence motifs and TFs that govern immune cell differentiation, strikingly conserved in human and mouse systems.

There is growing interest in applying deep learning computation to predict transcriptional activity and RNA processing from nucleotide sequences ^6,39,40^. A major breakthrough in using CNNs to accurately predict “activity profile” from sequence, which AI-TAC also benefits from, has been the utilization of multi-tasking frameworks that model multiple prediction tasks at once (e.g., predictions of accessibility across multiple tissues or cell types). The multi-task models can learn generalizable features whose combinations are predictive of different but related outcomes; this attribute is especially powerful in regulatory biology, where *combinations* of a finite set of sequence motifs underlies cellular differentiation. However, the representation of training data and the criteria for providing feedback to the model during the learning phase are of key importance, on which AI-TAC differs from previous work. By modeling *continuous* accessibility values across 81 cell populations that represent fine scale differences in immunocyte differentiation, and then measuring the model’s prediction error based on Pearson correlation, AI-TAC parameters were optimized to identify sequence features that are predictive of differences in profiles, rather than ubiquitous activity levels, a feature that proved essential to its performance. Our study also differs from previous work by its emphasis in robust extraction of learned motifs and its validation with epigenomic data.

Several elements may contribute to the excellent fit of the predicted profiles and motifs. Deep CNNs have outcompeted previous methods for the prediction of TF binding because the hierarchical nature of the model allows it to simultaneously learn both specific local patterns and long-range interactions in the sequence. Moreover, these models do not require manual curation of the input features enabling the identification of new important sequence features by the model. Although several recent works have introduced methods for dealing with longer input sequences such as dilated convolutions ^13,41,42^, the relatively short sequences that result from the ATAC-seq dataset precisely encapsulate regulatory regions, making the basic deep CNN model highly effective here.

The tight fit between AI-TAC’s “interpretation” and biochemical data gave high confidence that they were valid projections of the true regulation of chromatin accessibility across immunocytes. Furthermore, the cell-specific influence of these filters recapitulated prior knowledge about cell-type specificity for several transcription factors (e.g. Pax5 and Ebf1 in B-cells, Eomes/Tbx21 in NK cells, Spi1 and Cebp in myeloid cells). Several novel observations are worth highlighting: (1) The results yield a *high-resolution ranked landscape* of chromatin regulation across the entire immune system. Even if these players were recognized, their dominance (**Fig. 3a**) was not necessarily appreciated: knockouts only identify the stage at which a TF becomes essential for further differentiation, potentially distinct from those involved in overall specification of cell-specific chromatin architecture. Less expected was the dominant influence of Eomes/Tbx21 filters for NK cells, or of NRs for ILC2/3, which proved quite different from T cells (even the most differentiated NKT or effector CD8s), contradicting the over-simplification that ILCs are basically TCR-less T cells. This unique influence of NRs in ILC2/3, partially shared with RORg+ Tregs and some γδT, might prompt the speculation that these TFs and OCRs are active in cells at the microbial interface; it is also possible that these ILCs are further differentiated than any other cells in the dataset, a stage at which the NR family becomes more prominent. (2) *T cells as a default pathway*? Dominantly influential controllers were identified for B, myeloid and ILCs, but no strong equivalent emerged for T cells (influenced more weakly by Lef1/Tcf7, Tcf3, Ets, Runx and Gata). Unless a dominant T-determining factor was missed by AI-TAC, the implication is that T cell differentiation follows a different strategy. One might speculate that T cells are a lineage adopted when other avenues are no longer possible (i.e. by having terminally extinguished Spi1 and Pax5), or that the plasticity and functional diversity of T cells require flexible control, not compatible with a dominant master regulator. (3) 21 *novel motifs* were identified by AI-TAC (**Fig. S5**). Some of the un-annotated filters may represent “half-sites”, perhaps mere building blocks used by the CNN ^43^ or perhaps biologically relevant half-sites as reported for NF-κb or NRSF ^20,44^. Others appeared like typical TF binding sites (short and continuous blocks of preferred bases), and may represent unrecognized TFs or alternative sites for known TFs (TFBS databases are known to be incomplete) and require further investigation. Also intriguing were the poorly reproducible filters, which typically recognized scattered conserved bases; their low individual influence and non-reproducibility in different training runs would suggest that they only represent noise, but we cannot rule out that they correspond to a different regulatory syntax, perhaps read by non-coding RNAs. (4) The *repeat structure* (no tandem repeats outside of NF-κb and Sp1, but pervasive motif repeats in different enhancers connected to the same gene) suggests that eukaryotic genes do exploit cooperative multimeric interactions by repeats of the same factor, but do so by recruiting several spaced OCRs rather than by locally dense tandems, a solution that may provide both transcriptional and evolutionary flexibility. (5) *Novel TF combinations*. Deeper insights of co-occurring TF motifs were gained from combinatorial predictions (**Fig. 5**), again strongly validated by biochemical data. Some associations were expected (e.g. Pax5 and Ebf1), and the combination of AI-TAC and ChIP-seq validation data revealed patterns of differential association in B cell stages, as well as factors with broadly distributed co-influence (Tcf3 and Ets). But AI-TAC also identified 493 significant interactions, many previously unreported, some encompassing annotated filters (e.g. filter9, associated with NF-κb, Runx and Ets).

The underlying logic in this work is that, by analyzing how a deep neural network can decipher the cis-regulatory code of immune cell differentiation, we can infer how the biological network in live cells actually *does*. Some caveats need to be stated, however. Choosing correlation to determine the loss function improved predictions for variably-active loci, but penalized predictions of ubiquitously active OCRs. Another caveat is that CNNs leverage repeated effects, and will fail to identify very specific TF combinations that act only on one or two genes that may nevertheless be functionally critical (e.g. the λ5 enhancer ^28^ or the fine interplay between Tbx21 and Eomes during effector T cell differentiation ^45^). Transcription factors have varying degrees of dependence on sequence-specific DNA-binding: none for TFs such as Aire ^46^, variable for others such at the estrogen receptor [strictly dependent on a canonical motif at some loci, coopted in a looser manner at others ^47^]. AI-TAC would clearly miss TFs that do not rely on specific binding. Similarly, some TFs are “opportunistic”, only binding to chromatin already made accessible by other factors; FoxP3 is in this category ^48^, and it is interesting that no TF of the Forkhead family was discovered by AI-TAC, suggesting that Forkhead family factors may not be pioneers in hematopoietic lineages cells as they are in mesenchymal cells ^49^. TFs whose binding specificity is very dependent on dimer formation or on cofactors might be difficult for AI-TAC to recognize, although it is interesting to note that it is able to ferret out motifs for NF-κb, a TF family notorious for its combinatorial specificity and tolerance to variation ^20^. Relatedly, two factors competing for the same motifs may be poorly resolved by AI-TAC (e.g. the motif bound by Bcl11a and Bcl11b, essential for myeloid and T development respectively ^50,51^, appears mainly influential in myeloid and B cells). Finally, AI-TAC cannot read the influence of other means of regulation like specific DNA methylation, and there should be potential in integrating multiple data modalities into CNNs to further improve performance.

In conclusion, integrating a comprehensive *cis*-regulatory atlas of chromatin and transcript data with deep learning approaches has revealed modalities and complex patterns of immune transcriptional regulators, and how cell and lineage specificity across the immune system arise from the DNA sequence and can be encompassed in a genetic regulation network. Although some blind spots remain, this draft regulatory roadmap should provide a foundation to graft additional layers of human- or machine-generated results, and a springboard for experimental exploration.

## ACKNOWLEDGEMENTS

We thank Drs E. Rothenberg, A. Stark, B. Kee, S. Ghosh and A. Kundaje for insightful discussions. This research was enabled in part by computing support provided by WestGrid (www.westgrid.ca) and Compute Canada (www.computecanada.ca). This work was supported by grants AI072073 from the NIH to ImmGen, NSERC DGP to SM^2^. SM^1^ was partially supported by an NSERC CREATE scholarship and KM was partially supported by NSERC USRA. RR was supported by NIH supplement 3R01AI116834-03S1.

## Data and Software Availability

All ATAC-Seq datasets from the ImmGen project are available from www.immgen.org and GEO (accession number GSE100738). All software and scripts for generating the AI-TAC model will be available from https://github.com/smaslova/AI-TAC.

