## Supplementary Methods and Figures for "Learning immune cell differentiation"

#### Methods and Development Notes

##### Tuning AI-TAC's architecture

In tuning the model's various hyperparameters, Bayesian optimization was performed using skopt [14] package to progressively search for the best set of parameters in the given search space. The following hyperparameters were optimized: 1) number of convolutional layers and fully connected layers, 2) filter size, pooling size, pooling stride and dilation rate for convolution layers, 3) number of filters for each layer, and 4) type of activation function for each layer. The number of search iterations was set to be 130, where in each iteration the model was trained and evaluated with a new set of hyperparameters. We ranked the models based on the correlation *loss* used in AI-TAC (measured as error = 1 - Pearson correlation coefficient).

We observed that models with similar hyperparameter setting as Basset [7] (specifically those with 3-4 convolutional layers, and 300-500 filters of size 13-21bp in the first layer) were among the top performing models (best observed loss=0.61, "Basset" loss=0.64). We also explicitly assessed hyperparameter settings similar to DeepSea [13] and DeeperBind [52] but observed that both of these model settings were less performant (loss=0.69 and loss=0.85, respectively).

Based on the results from Bayesian Optimization analysis, we opted to create a CNN ("AI-TAC") with similar architecture as the Basset model which includes three convolutional layers followed by two fully connected layers (**Fig. S1a**). In the case of AI-TAC, the first layer takes as input a 251x4 matrix (one-hot-encoding of the 251bp DNA sequence underlying a given OCR). The final output layer consists of 81 neurons, corresponding to chromatin accessibility profile across the 81 immune populations measured.

The first three convolutional layers utilize the ReLU activation function, followed by maxpooling and batch normalization, with the very first layer consisting of 300 19bp filters. The two fully connected layers each consists of 1000 neurons and also use the ReLU activation function. During training, a dropout rate of 0.03 was applied to the fully connected layers as a form of regularization. The model was implemented in PyTorch version 0.4.0. The training was performed using the Adam optimizer [53] for 10 epochs with a learning rate of 0.001 and a mini-batch size of 100.

**Training AI-TAC and measuring it performance**

The ImmGen ATAC dataset consists of 327,927 high-confidence OCRs from the mouse genome (excluding the X and Y chromosomes) that were previously described [2]. Each OCR's accessibility profile across 81 immune populations was quantified as its quantile normalized log count value. The AI-TAC model was trained on a randomly selected 90% of the OCRs in our dataset, and the remaining 10% was retained as the test set.

We assessed the robustness of the model's performance in several ways. First, to ensure that the test set accuracy obtained by our model is reproducible, we performed 10-fold cross-validation (CV) on the entire dataset ten individual times (yielding 100 trained models). In this way, each of the 327,927 OCRs was considered as a test OCR by ten different trained models (**Fig. S2c**). Encouragingly, we observed stable performance for well-predicted OCRs (**Fig. S2d**), suggesting that training examples are sufficiently representative.

Second, because of the possible repeated representation of transcription factor binding sites (TFBS) within a short DNA segment, we also assessed the model's performance with a "leave-out-chromosome" approach. Specifically, we trained 19 models, each leaving out a different mouse autosome. We observed similar test set performance with the chromosome-leave-out-approach as the random 10% leave out approach (**Fig. S2b and S2c**).

Third, we compared the performance of our model on "real data" to three different approaches to randomization of the relationship between OCRs and their activity:

1. Randomly shuffled 251 base pair input sequences
2. Randomly shuffled ATAC-Seq activity vectors of length 81
3. Randomly permuted order of input sequence and output vectors in the dataset

As shown in **Figure S2a** and **Figure 1c**, we observed that randomization of the relationship between OCR sequence and activity profiles yielded the expected null, indicating "real" signal and structure in the dataset that the model is able to exploit to make predictions.

**Deriving motifs by analyzing first layer filters**

The 300 first layer convolutional filters were analyzed by converting them to position weight matrices (PWMs). To do so, for each first layer filter, we first identified all 19bp sequences that activate the filter by at least ½ of the maximum activation for that filter across all well-predicted (correlation >0.75) OCRs [7]. Next, we constructed a position frequency matrix based on the prevalence of each of nucleotide along the 19bp long sequences, and finally we converted the position frequency matrix to a PWM by using a background uniform nucleotide frequency of 0.25. This analysis yielded 300 PWMs, each capturing the motif that is detected by a first layer filter.

To understand how reproducible the learned first layer filters are we trained 10 additional models using different 90% subsets of the data, then extracted the 300 filter PWMs from all 10 models. We measured the similarity between the filters of AI-TAC and those from each of the other models using the TomTom PWM comparison tool [18]. We then defined a reproducibility score for each of the AI-TAC filter motifs as the number of models with at least one matching motif using an FDR q-value cut-off of 0.05 on the TomTom results.

**Computing filter influence**

The impact of each filter was computed by effectively removing the filter from AI-TAC and quantifying the impact on the model's prediction. Specifically, we replaced all activation values for the given filter with its average activation value across all samples in the batch, then fed the output through the remaining layers of the model to obtain the altered prediction vector. Based on this nullification process, we computed two quantities. First, the *overall influence* value for a given filter was computed as the average (across OCRs) of the squared difference between the correlation in prediction of accessibility profiles (loss) of the altered and un-altered model.

Second, to understand the cell-type-specific impact of each filter we also computed an *influence profile* per filter, as the average (across OCRs) of the squared difference between the original and modified prediction values for each output neuron (corresponding to a given cell population). We additionally computed a “signed” version of the influence profile, by taking the difference between altered and un-altered prediction for each neuron, which shows whether the presence of each filter is predictive of higher or lower chromatin accessibility in each cell population (**Fig3a**).

All influence values were computed using 51,732 OCRs for which the AI-TAC prediction has greater than 0.75 correlation with the ground-truth chromatin accessibility. To ensure that including well-predicted OCRs from the training set would not bias our results, we compared the influence values obtained from well-predicted test and training set OCRs (**Fig. S2f)**. The influence values computed on these two sets were strongly correlated. Additionally, the filter PWMs from the same model obtained from the test versus training set OCRs were virtually identical, with a TomTom q-value of $2.9 \times 10^{-27}$ or less for all but one pair of test set and training set derived PWMs.

**Assigning TFs to filters**

The AI-TAC first layer filter motifs were compared to the Cis-BP *Mus Musculus* database version 1.02 of known transcription factor motifs using TomTom [17]. Results are provided in **Table S2**.

##### Assigning TFs to filters based on analysis of RNA-seq gene expression data

For each filter, we computed its accessibility profile as the average accessibility of the top 500 OCRs activated by that filter. We then computed the correlation between this accessibility profile and expression levels of each eligible TF match for that filter (**Table S3**). For this analysis, we only considered TF matches for each filter whose Cis-BP q-values were within 2-orders of magnitude of the lowest observed Cis-BP q-value.

##### Fine-tuning AI-TAC with 99 first layer filters

To understand how much of the model performance can be recovered with only the 99 filters reproduced in 80% of our models we tested a version of AI-TAC with all other first layer filters removed. A new model with 99 first layer filters was initialized with the reproducible weights from AI-TAC and train for 10 additional epochs and tested the model at the end of each epoch on a validation set. The best performing model was found after just one epoch of fine-tuning. We then computed the prediction correlation values on the test set OCRs (10% of the data) for both the original and fine-tuned AI-TAC models (**Fig S2e**). The prediction correlation values were significantly different for only 10% of OCRs (using FDR-corrected p-value cut-off of 0.05).

##### Comparing first layer filter motifs to DeepLIFT-derived motifs

To ensure that we were not missing motifs important to the model that were represented partially in the first layer and assembled in deeper layers [43] we tried using an alternative method of deep neural network interpretation, DeepLIFT [15]. Rather than extracting the first layer only, DeepLIFT computes a score for each position of an input sequence by backpropagating the contribution of the existing base pair to each of the output neurons.

To apply DeepLIFT to our model we first re-trained the AI-TAC architecture with an additional sigmoid activation on the output layer in order to bound the values of predicted chromatin activity for each cell type between 0 and 1. We selected all test set OCRs for which the prediction correlation was greater than 0.75 (2305 sequences in total) and applied DeepLIFT (v. 0.6.7.1) with the genomics default settings and 10 reference sequences per sample to compute the attribution scores of each base pair to the model output.

To compile the DeepLIFT we used TF-MoDISco [15] which clusters short sub-sequences with high DeepLIFT attribution scores into motifs. We applied TF-MoDISco (v. 0.4.2.3) with an initial window of size 10 (sliding_window_size) and an FDR value of 0.05 (target_seqlet_fdr) to identify "seqlets" with high attribution scores. The windows are then expanded by 5 bp on either side (flank_size), and the resulting sequences are clustered by similarity. Any cluster with less than 60 seqlets is broken up and re-assigned to other clusters, and the final motifs are then expanded to 45 base pairs.

A total of 424 motifs were identified using DeepLIFT and TF-MoDISco, and for each of these we computed one score by averaging the DeepLIFT attribution score from all sequences contributing to the motif and all positions across the motif. We then compared all DeepLIFT-generated motifs to the AI-TAC first layer filter motifs using TomTom (Figure S3A-B). This comparison showed that DeepLIFT recovers some, but not all, important reproducible motifs from AI-TAC (Fig. S3a), and does not appear to discover any additional motifs that weren't identified by AI-TAC (Fig. S3b).We additionally computed an approximate motif length as the distance between the first and last base pair with an information content of 0.2 or more. Although the DeepLIFT motifs were expanded to 45 base pairs, we found the vast majority of them were still shorter than 19 bp (Figure S3C); one of the two exceptions is shown in Figure S3D and appears to correspond to the GC content of the sequence.

##### Transcription factor ChIP-seq data processing

ChIP-seq raw data were retrieved for Pax5 from *in vivo* proB and mature B cells [[27], GSE38046], Ebf1 from Pro B cells [[27], GSE78844], Tcf1 (referred to as Tcf7) from T.DP [[31], GSE46662], and Spi1 (Pu.1) [[54], GSE62826] in macrophages. Fastq files were mapped to the mm10 reference genome using bowtie2 [55] and samtools [56]. Duplicate reads were removed using Picard (http://broadinstitute.github.io/picard), bigWig tracks were generated using deepTools2 [57] and visualized with IGV [58]. TF ChIP-seq peaks were called for Pax5, Ebf1, and Tcf7, Spi1 using the HOMER [59] *findPeaks* function with an FDR of 1% using the parameter (*-style factor),* also including respective input background determined peaks for each respective ChIP-seq dataset. Intersection of TF peaks with corresponding replicates was performed using the BEDtools [60] *intersect* function with a 50% reciprocal overlap requirement, resulting in 9,655 Ebf1, 45,348 Pro B Pax5, 28,009 Mature B Pax5, 22,000 Spi1 and 14,042 Tcf7 peaks.

##### Deriving Pax5 footprint

Centering on filter 167 (Pax5) and 275 (Ctcf) sequences, Tn5 insertions were mapped in order to extract the AI-TAC Pax5 footprint trace. To validate the AI-TAC derived footprint, we used Pro B cell Pax5 ChIP-seq peaks to first identify a Pax5 footprint *de novo* using a hidden Markov model approach HINT-ATAC [23]. ATAC-seq footprints were derived from Pax5 binding sites (*rgt-hint footprinting --atac-seq --paired-end*), matched to motifs (*rgt-motif analysis matching)* and used to resolve a Pax5 footprint trace based on B cell Tn5 insertions (*rgt-hint differential –bc).*

**Validation of AI-TAC predictions with immune transcription factor ChIP-seq**

We validated predictions for filters 167 (Pax5), 260 (Ebf1) and 166 (Lef1/Tcf7) using *in vivo* ChIP-seq data for these TFs. We estimated the proportion of top 500 OCRs (ranked by influence) that were supported by a TF ChIP-seq peak respectively. Filter 167 (Pax5) was used as a negative control for validation of filter 166 (Lef1/Tcf7) sequences, and performed inversely for validation of filters 260 and 167. A global validation and comparison was performed for Pax5 binding from Mature B and Pro B cells for OCRs (top 500) across all 99 reproducible filters. In a similar manner, comparisons between Tcf7 and Pax5 were performed for OCRs (top 500) across all 99 reproducible filters. To further investigate OCRs with shared Pax5 and Ebf1 filter influences, we first computed OCR intensities for predicted OCRs using the maximal OCR intensity (normalized ATAC-seq counts) for a respective lineage and subtracting the maximal value across all cell types in all other lineages. Of the predicted OCRs (40,183), 5,443 OCRs demonstrate B cell specific activity. We randomly selected 5,443 non-B cell OCRs as a comparison from the remaining set. Pax5 and Ebf1 ChIP-seq intensities were mapped to predicted B cell specific and non-B cell OCRs.

##### Identifying co-influential filter pairs

To identify pairs of motifs that are co-influential, we first assigned a set of influential filters to each well predicted OCR by using overall influence threshold of 0.0025 (which corresponds to >5% change in correlation for prediction of an OCRs profile as the result of that filter's nullification). In identifying pairs that co-occur as influential, we computed a hypergeometric p-value quantifying the significance of co-occurrences as compared to expectation based on filter prevalence (**Table S5**). Because filters that are redundant with each other (e.g., two versions of Pax5 motifs) can be counted as co-influential, we removed from consideration pairs of filters whose PWM were similar to each other at PWMEnrich (https://www.bioconductor.org/packages/release/bioc/html/PWMEnrich.html) value of >0.5.

##### Filter co-influence analyses of AI-TAC's fully-connected layer

Of the 1000 nodes with activation in AI-TAC's last fully-connected layer before the output layer, 695 were assessed for filter co-influential patterns of predicted OCRs (n=30,875). A two-dimensional representation and clustering of OCR patterns in the last hidden layer was performed [61]. Lineage specificity across major lineages was computed as follows: (1) the max intensity for each OCR was defined across all cell-types grouped as lineages; (2) For a given lineage, the max ATAC intensity was subtracted for the respective lineage from the max of all other compared lineages. Single and co-filter patterns were mapped based on the filter influence threshold (0.0025).

##### Clustering of reproducible AI-TAC filters

Clustering of all 99 reproducible filters was performed using RSAT [62]. Briefly, filters were clustered using the matrix-clustering function with a Pearson correlation of ≥0.7 for identifying filter clusters, represented in dendrogram or heatmap formats.

##### Application of AI-TAC to human data

ATAC-seq from 18 human cell types were downloaded from [5], and processed using the same parameters used in the ImmGen ATAC project [2] (i.e., MACS2 algorithm for peak calling to obtain 251bp peaks). In total, ATAC intensities for 539,611 OCRs passed the threshold of > 3 reads, and these were used for further analysis. As a first analysis, AI-TAC model trained in mouse was directly applied to these human OCRs. 8 of the human cell types matched to those predicted from mouse data, mostly at lineage level (**Table S7**). When appropriate, the predictions were averaged for multiple cell populations to represent the predictions at the lineage level.

We then explored fine-tuning AI-TAC model on human dataset. We adopted the same network architecture as used in the mouse-trained model, except that the last output layer was truncated and replaced with a layer of 18 output neurons which match the number of cell types in the human dataset. Using the mouse-trained model weights as the initial training state, we applied a 0.0001 learning rate (0.1 times of the original one) to retrain the model weights on human data for four epochs decided by an early stopping strategy.

To compare the mouse and human trained model, we used 50% of human OCRs to train an equivalent CNN model (same architecture as that used in mouse), but only on the 8 of human cell types which are matched to the mouse cell types. For the remaining 50% OCRs as the test set, the predictions of the human model were compared to that of AI-TAC (**Fig. 7c**).

#### SUPPLEMENTARY FIGURE LEGENDS

**Fig. S1:** **a**, The architecture of the AI-TAC model. **b**, Results of the Bayesian Optimization: each dot depicts a model specification (particular setting of hyperparameters), y-axis depicts the total loss/error (1-correlation) on test data. **c**, Test set prediction correlation for a model trained with mean squared error loss and Pearson correlation loss (center line, median; box limits, upper and lower quartiles; whiskers, 1.5x interquartile range; points, outliers). The data is split by quantiles of the coefficient of variation values of the ATAC-Seq signal of each OCR. **d**, Coefficient of variation of chromatin activity (x-axis) versus prediction correlation (y-axis) of test-set OCRs, from a model trained with mean squared error loss (left) and Pearson correlation loss (right).

**Fig. S2:** **a**, Performance of the model (measured by Pearson correlation) on real and shuffled dataset. (Top) randomization by shuffling the sequences, and (Bottom) by shuffling the assignment of each OCR to its accessibility profile. For randomization experiments, models were trained and tested on shuffled data. **b**, Boxplot showing the test performance of 19 separate models, trained by leaving each of the 19 autosomes out once (center line, median; box limits, upper and lower quartiles; whiskers, 1.5x interquartile range; points, outliers). **c**, In each trial, 10 models were trained using 10-fold cross validation (CV) using a random partition of datasets. Based on the 10 trials of 10-CV experiments, each OCR was considered a test OCR in 10 different models. Boxplot shows the test error for all OCRs for each 10-fold cross-validation trial. **d**,

Figure shows the mean prediction correlation for each test OCR across the 10 independently trained models. Y-axis shows the range of prediction correlation for each OCR. **e**, 2D histogram plot of the prediction correlation values of the full AI-TAC model versus fine-tuned AI-TAC with 99 reproducible motifs only, computed over OCRs not used for fine-tuning. **f**, Scatter plot of influence values per filter when computed on well-predicted OCRs in test set only (x-axis), versus its computation from well-predicted training set OCRs (y-axis).

**Fig. S3:** **a**, For each of the 300 AI-TAC filters, the negative q-value for the best matching DeepLIFT generated motif plotted against the influence of each filter. Colored by the reproducibility metric of each filter motif (the number of training runs in which that filter was learned, out of 11.) Significant q-value of 0.01 indicated with blue line. **b**, The average DeepLIFT score of all motifs discovered by the DeepLIFT/TF-MoDISco pipeline versus the TomTom q-value for the best matching AI-TAC filter motif. Colored by the reproducibility value of the best matching AI-TAC filter motif. **c**, Histogram of DeepLIFT motif lengths. **d**, PWM of the longest motif (length=40) discovered by DeepLIFT.

**Fig. S4:** **a**, An example showing the correlation between RNA-seq expression of a TF (Pax5) versus the average accessibility profile of OCRs that are activated by the corresponding filter (167) across 81 cell-types. The correlation coefficient of expression versus accessibility profile is 0.97. **b**, For each filter PWM, the eligible TF matches based on CIS-BP results were considered in correlation analysis, where the correlation between gene expression profile of the TF and accessibility profile of the activated

OCRs across 81 cell populations was computed. For each filter, the correlation coefficients for eligible TFs are shown on the y-axis. Colors represent the influence values for each filter. **c**, Figure shows examples of PWMs were multiple redundant representation of the same TF was found for Lef1/Tcf7 (50 and 166) Ctcf (23 and 69), Irf1 (165 and 250) and NfkB variants (231, 247 and 65). **d**, Filter clustering (Pearson 0.6) of Nfkb filters and motif variants across AI-TAC runs.

**Fig. S5:** Reproducible filters that did not match to any Cis-BP TF motifs with q-value < 0.05 and did not get assigned to any TF matches manually. The lineage most influenced by each filter is listed, defined as the lineage with the highest mean cell-specific influence.

**Fig. S6:** **a**, The AI-TAC model was fine-tuned for prediction of T cell populations, by retraining the model for one epoch. Figure shows the influence values of all filter motifs/TFs across T cell populations. **b**, Observed and AI-TAC predicted activity comparing Tregs and Macrophages. Top 2000 OCRs bound by Pu.1 ChIP-seq are colored. **c**, Observed and AI-TAC predicted activity comparing NkT and NkT with LPS treatment after 3hrs. OCRs with influence (0.0025) for 231 (NF-$\kappa$b-het, n=311) are colored.

**Fig. S7:** **a**, The maximum weight between a subset of reproducible first layer filters second layer convolutional filters. All first layer filters belonging to an RSAT cluster with more than one motif are shown. The second layer filters were selected based on a weight threshold of 0.7 for at least one of the first layer filters. For a few examples, we visualized second layer filter weights for the most heavily weighted first-layer motifs along with the corresponding first layer motif PWMs. **b,c**, The figure shows two instances of a second layer motif aggregating similar first layer motifs. **d,e**, Two examples of a second layer filter recognizing first layer motifs that are reverse compliments.

**Fig. S8:** Histogram of number of regulators assigned to each gene based on the inferred regulatory network.

Figure S1

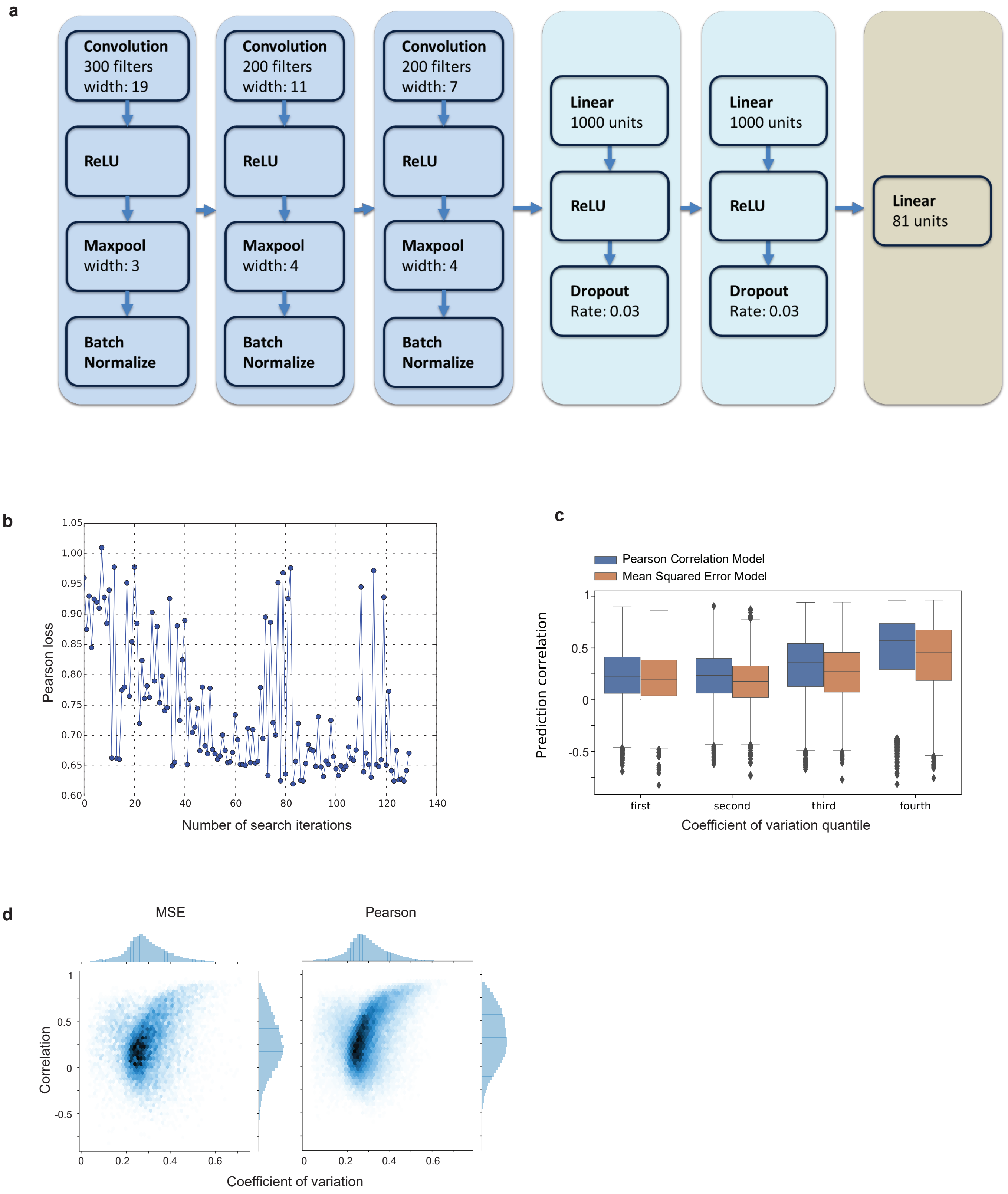
a
Convolution
300 filters
width: 19
ReLU
Maxpool
width: 3
Batch
Normalize
Convolution
200 filters
width: 11
ReLU
Maxpool
width: 4
Batch
Normalize
Convolution
200 filters
width: 7
ReLU
Maxpool
width: 4
Batch
Normalize
Linear
1000 units
ReLU
Dropout
Rate: 0.03
Linear
1000 units
ReLU
Dropout
Rate: 0.03
Linear
81 units
b
Pearson loss
Number of search iterations
c
Pearson Correlation Model
Mean Squared Error Model
Prediction correlation
first
second
third
fourth
Coefficient of variation quantile
d
MSE
Pearson
Correlation
Coefficient of variation

Figure S2

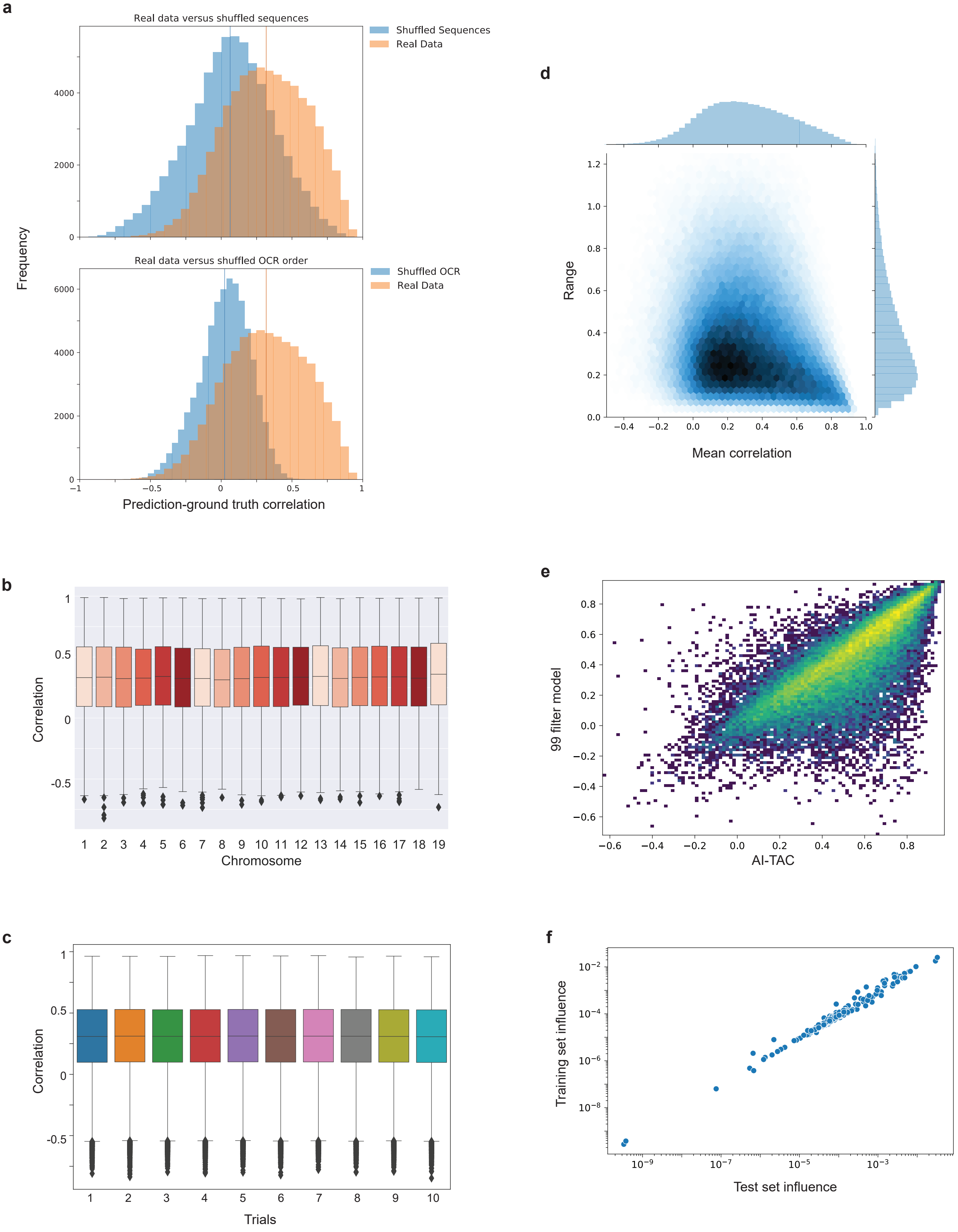
a
Real data versus shuffled sequences
Shuffled Sequences
Real Data
Real data versus shuffled OCR order
Shuffled OCR
Real Data
Frequency
Prediction-ground truth correlation
b
Correlation
Chromosome
c
Correlation
Trials
d
Range
Mean correlation
e
99 filter model
AI-TAC
f
Training set influence
Test set influence

Figure S3

a

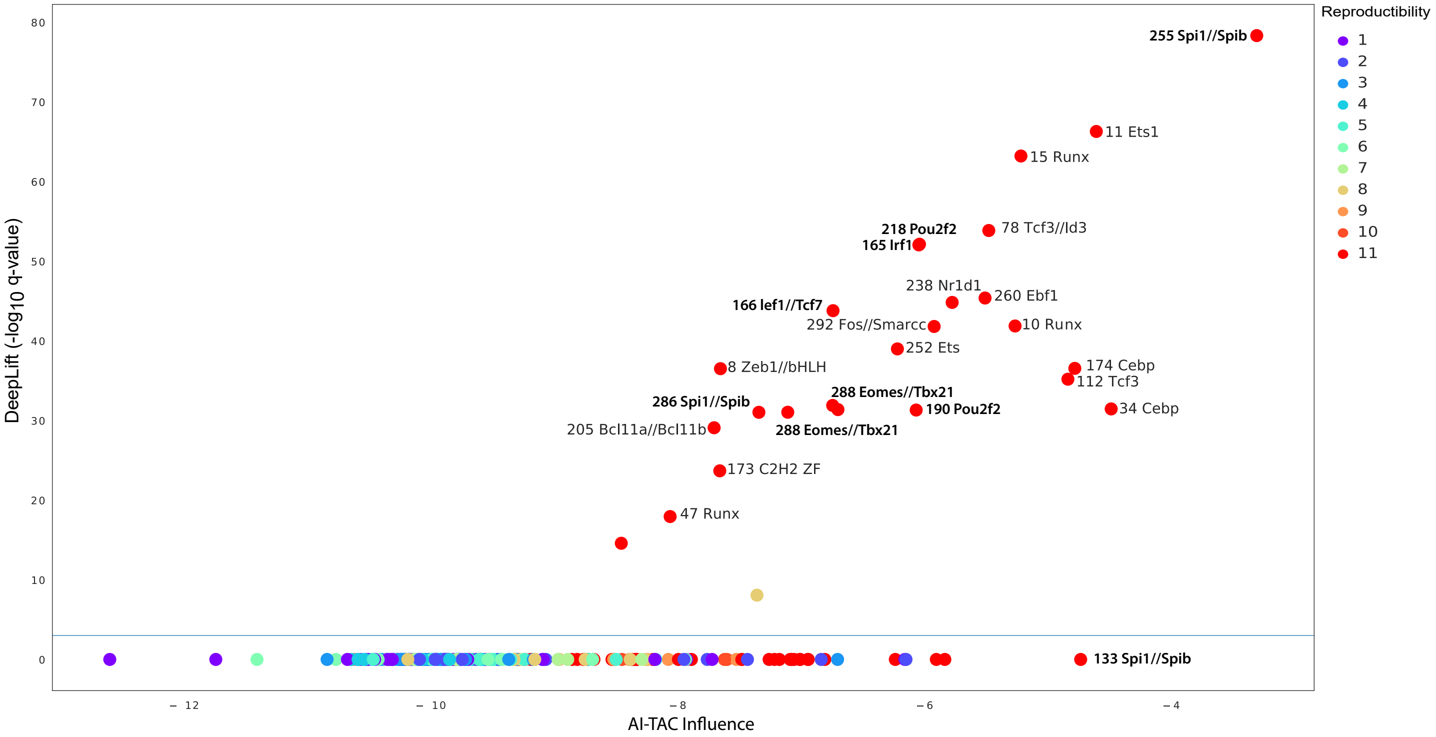


b

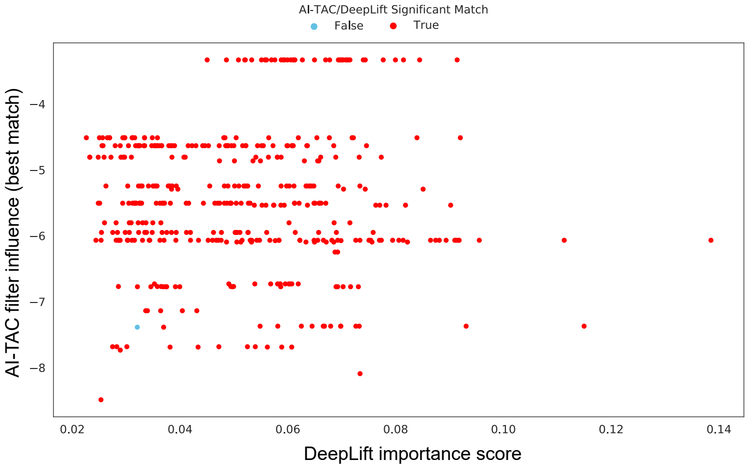


c

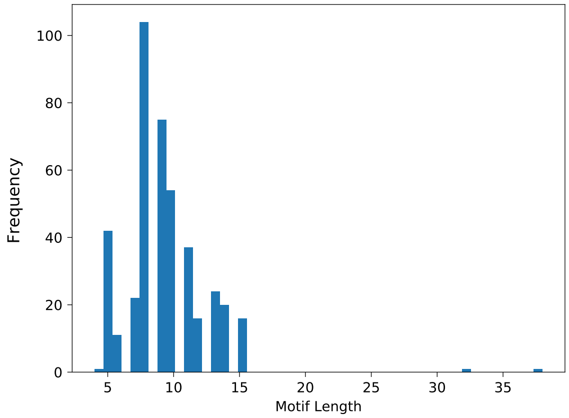


d

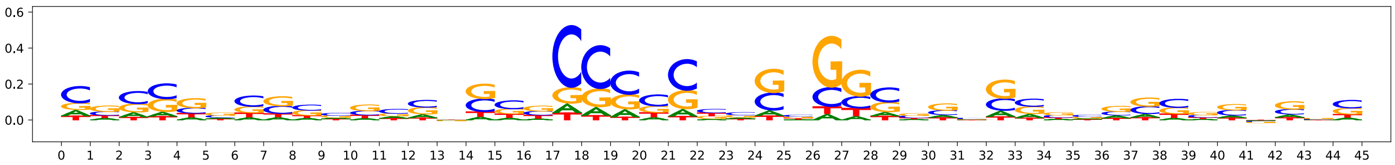

### Figure S4

a

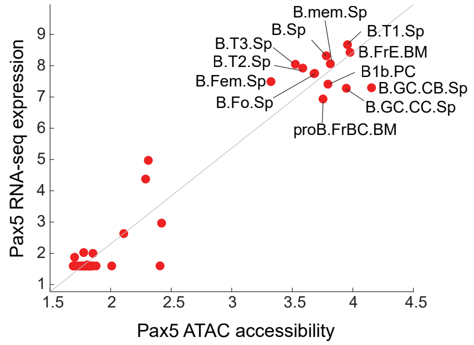


b

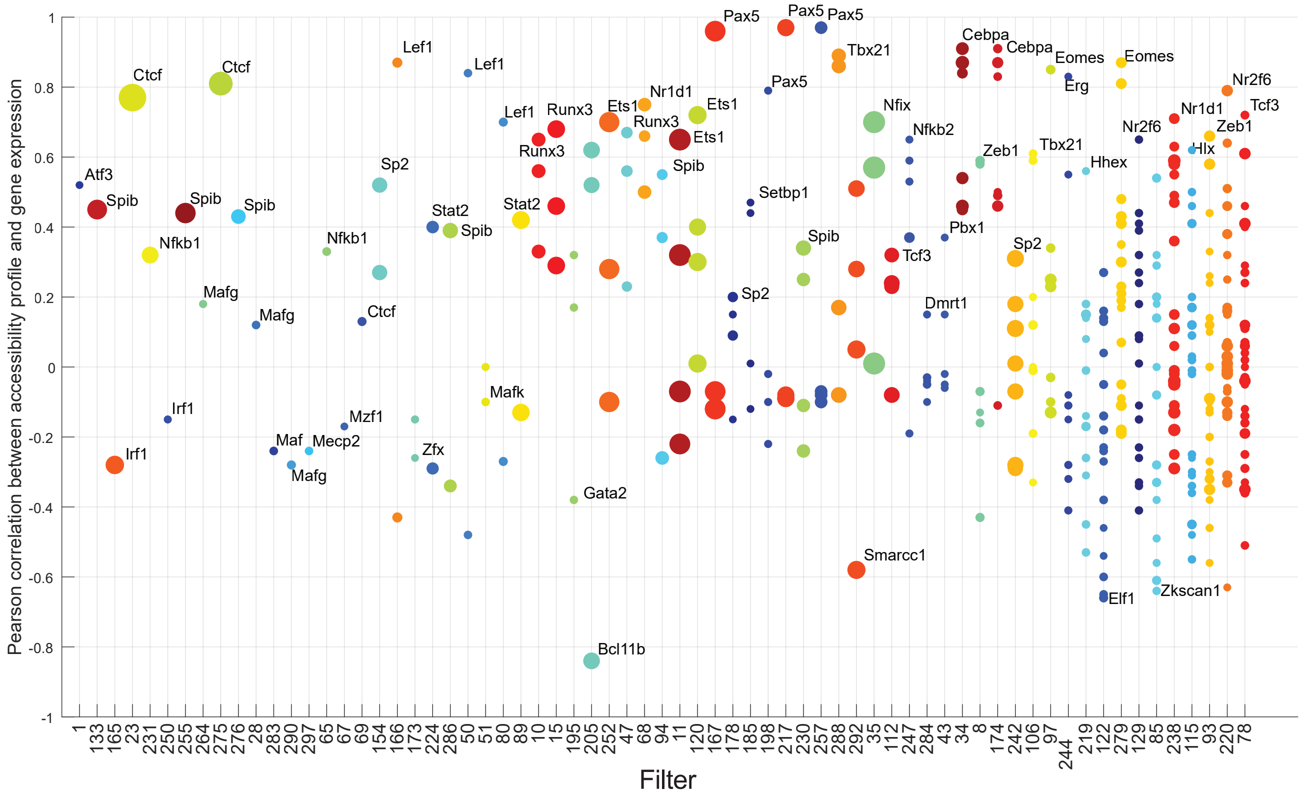


c

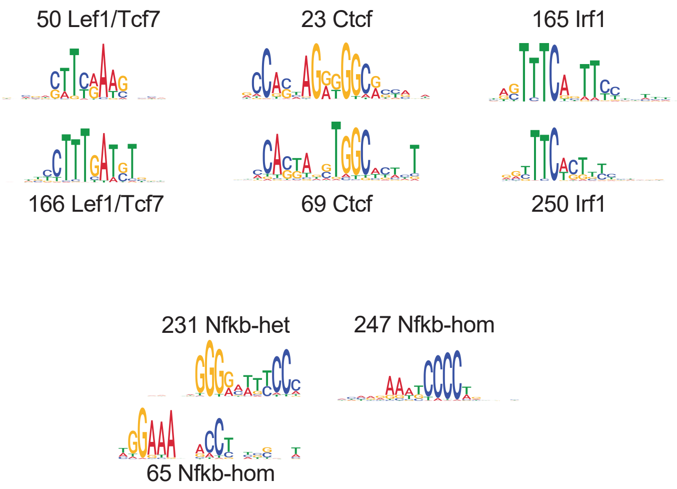


d

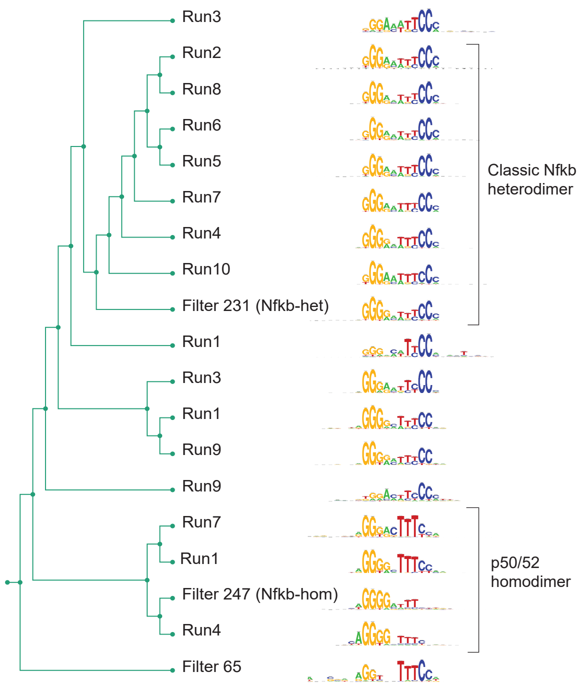

Figure S5

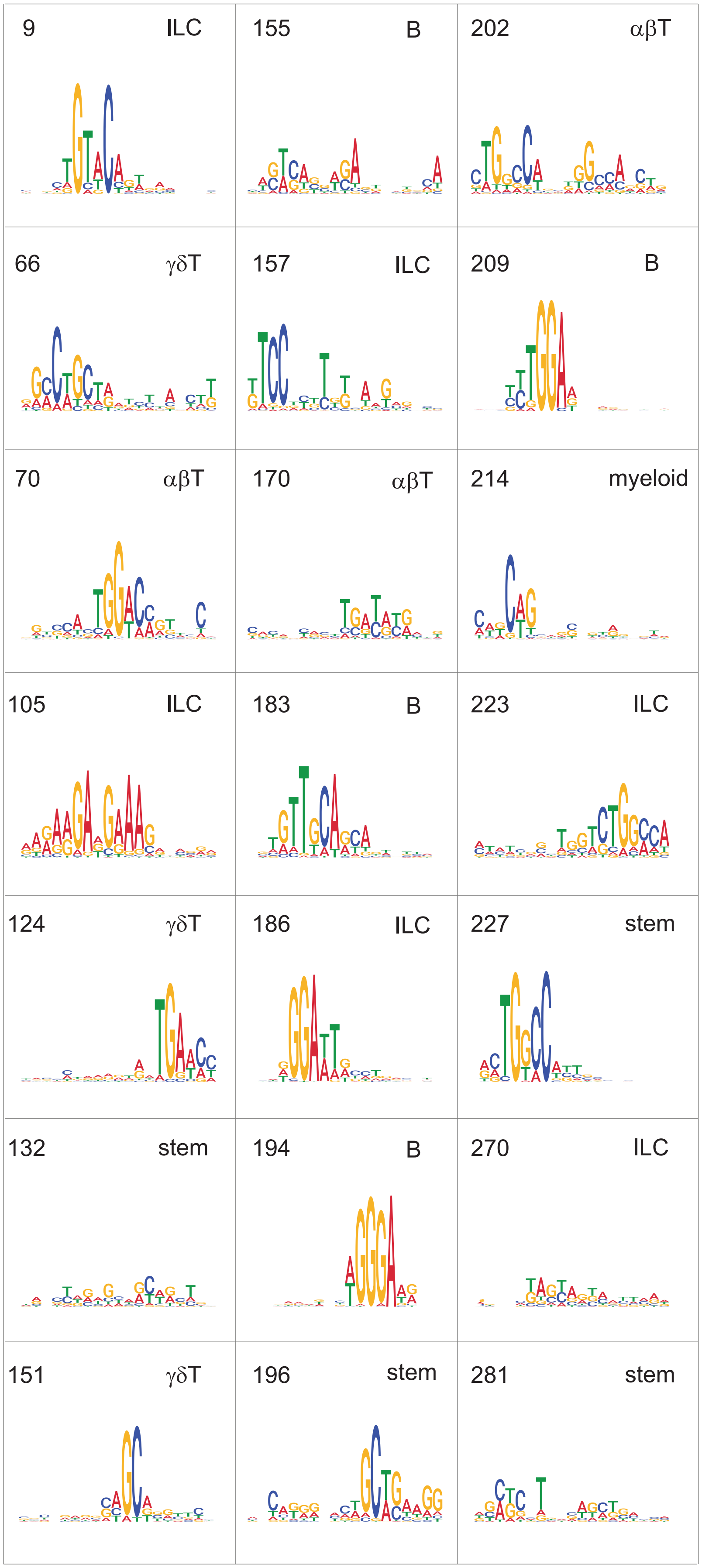
9 ILC
155 B
202 αβT
66 γδT
157 ILC
209 B
70 αβT
170 αβT
214 myeloid
105 ILC
183 B
223 ILC
124 γδT
186 ILC
227 stem
132 stem
194 B
270 ILC
151 γδT
196 stem
281 stem

### Figure S6

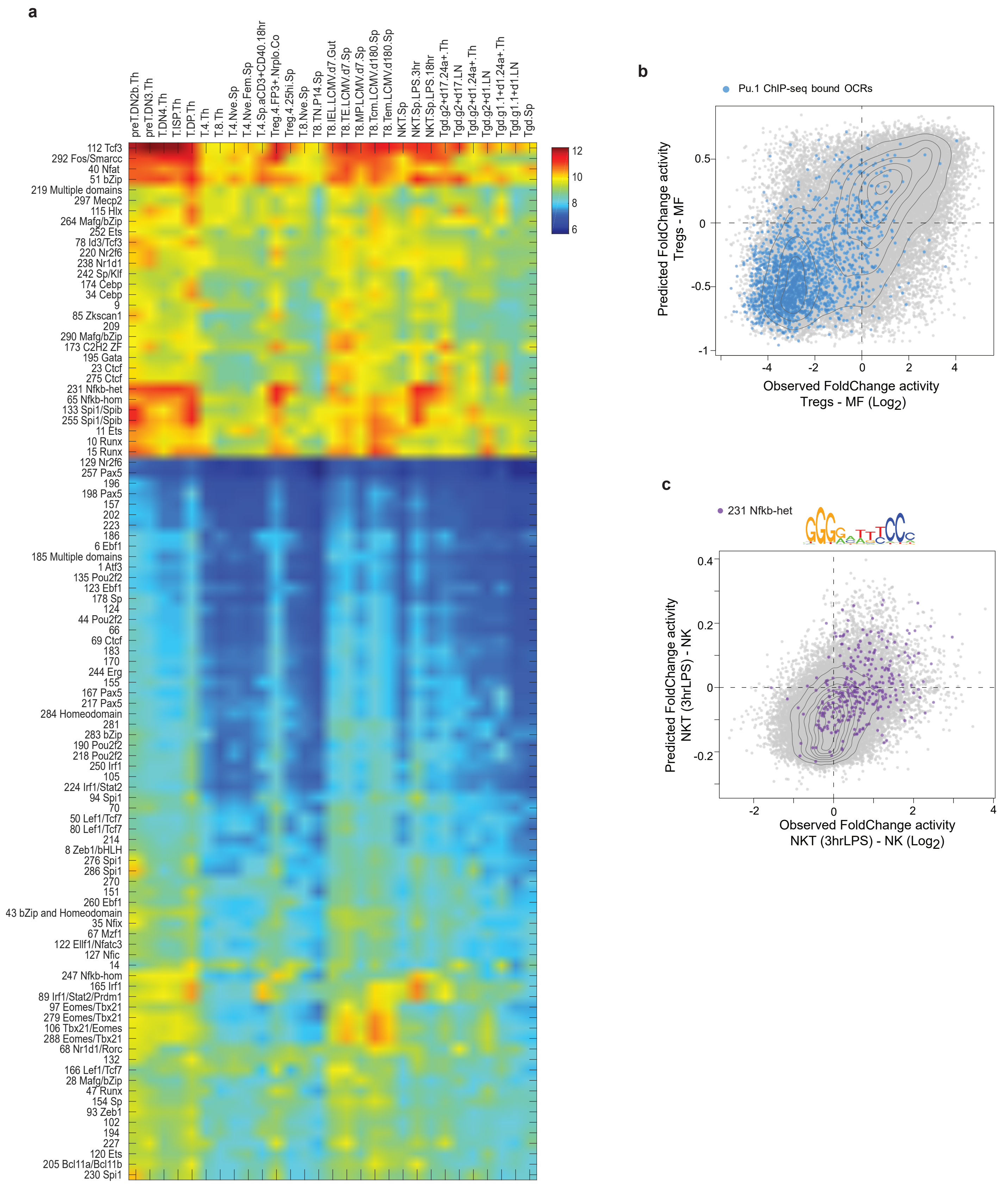
a
preT.DN2b.Th
preT.DN3.Th
T.DN4.Th
T.ISP.Th
T.DP.Th
T.4.Th
T.8.Th
T.4.Nve.Sp
T.4.Nve.Fem.Sp
T.4.Sp.aCD3+CD40.18hr
Treg.4.FP3+.Nrplo.Co
Treg.4.25hi.Sp
T.8.Nve.Sp
T8.TN.P14.Sp
T8.IEL.LCMV.d7.Gut
T8.TE.LCMV.d7.Sp
T8.MP.LCMV.d7.Sp
T8.Tcm.LCMV.d180.Sp
T8.Tem.LCMV.d180.Sp
NKT.Sp
NKT.Sp.LPS.3hr
NKT.Sp.LPS.18hr
Tgd.g2+d17.24a+.Th
Tgd.g2+d17.LN
Tgd.g2+d1.24a+.Th
Tgd.g2+d1.LN
Tgd.g1.1+d1.24a+.Th
Tgd.g1.1+d1.LN
Tgd.Sp
112 Tcf3
292 Fos/Smarcc
40 Nfat
51 bZip
219 Multiple domains
297 Mecp2
115 Hlx
264 Mafg/bZip
252 Ets
78 Id3/Tcf3
220 Nr2f6
238 Nr1d1
242 Sp/Klf
174 Cebp
34 Cebp
9
85 Zkscan1
209
290 Mafg/bZip
173 C2H2 ZF
195 Gata
23 Ctcf
275 Ctcf
231 Nfkb-het
65 Nfkb-hom
133 Spi1/Spib
255 Spi1/Spib
11 Ets
10 Runx
15 Runx
129 Nr2f6
257 Pax5
196
198 Pax5
157
202
223
186
6 Ebf1
185 Multiple domains
1 Atf3
135 Pou2f2
123 Ebf1
178 Sp
124
44 Pou2f2
66
69 Ctcf
183
170
244 Erg
155
167 Pax5
217 Pax5
284 Homeodomain
281
283 bZip
190 Pou2f2
218 Pou2f2
250 Irf1
105
224 Irf1/Stat2
94 Spi1
70
50 Lef1/Tcf7
80 Lef1/Tcf7
214
8 Zeb1/bHLH
276 Spi1
286 Spi1
270
151
260 Ebf1
43 bZip and Homeodomain
35 Nfix
67 Mzf1
122 Ellf1/Nfatc3
127 Nfic
14
247 Nfkb-hom
165 Irf1
89 Irf1/Stat2/Prdm1
97 Eomes/Tbx21
279 Eomes/Tbx21
106 Tbx21/Eomes
288 Eomes/Tbx21
68 Nr1d1/Rorc
132
166 Lef1/Tcf7
28 Mafg/bZip
47 Runx
154 Sp
93 Zeb1
102
194
227
120 Ets
205 Bcl11a/Bcl11b
230 Spi1
12
10
8
6
b
Pu.1 ChIP-seq bound OCRs
Predicted FoldChange activity
Tregs - MF
0.5
0
-0.5
-1
-4
-2
0
2
4
Observed FoldChange activity
Tregs - MF (Log2)
c
231 Nfkb-het
0.4
0.2
0
-0.2
-2
0
2
4
Predicted FoldChange activity
NKT (3hrLPS) - NK
Observed FoldChange activity
NKT (3hrLPS) - NK (Log2)

Figure S7

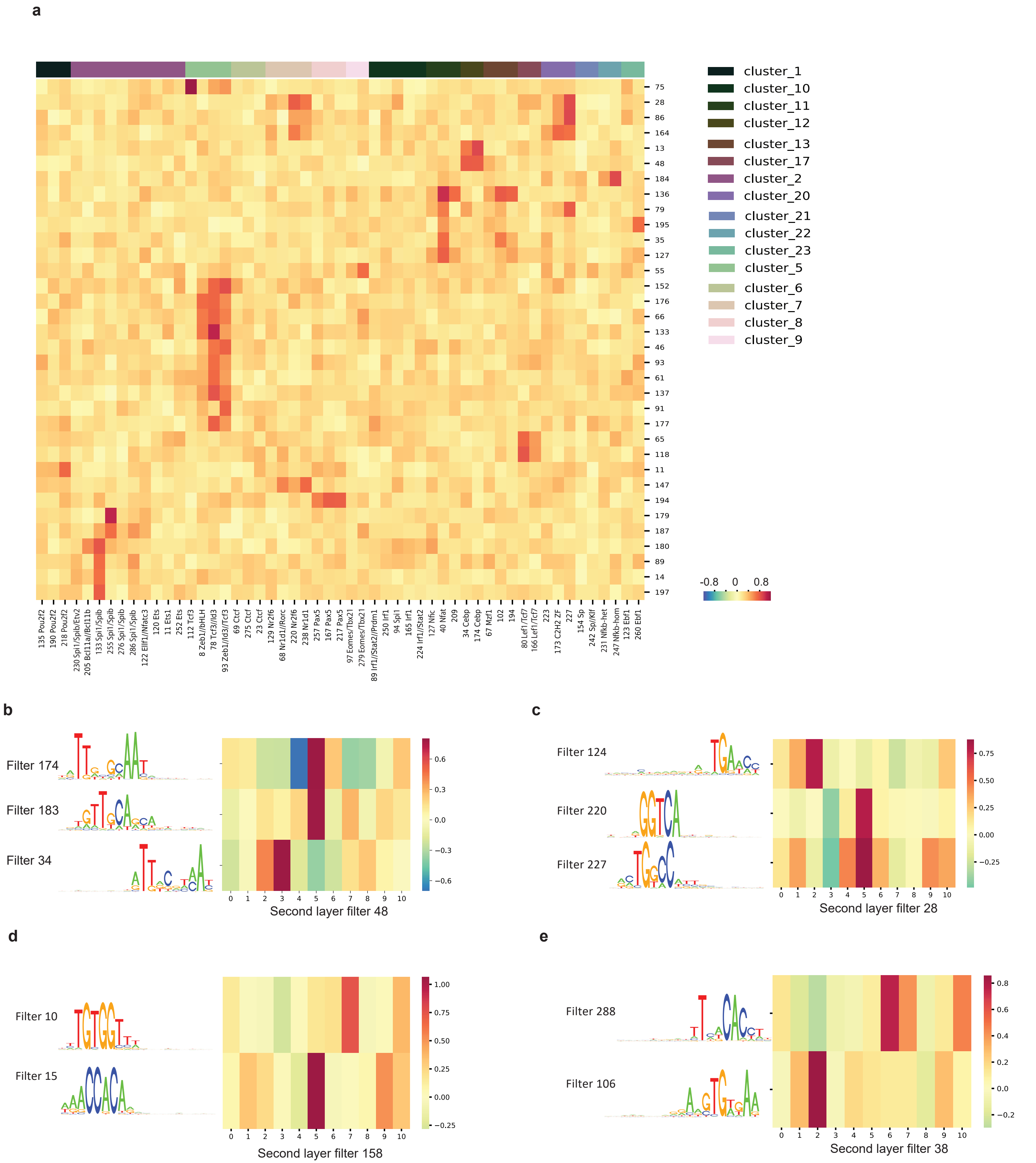
a
cluster_1
cluster_10
cluster_11
cluster_12
cluster_13
cluster_17
cluster_2
cluster_20
cluster_21
cluster_22
cluster_23
cluster_5
cluster_6
cluster_7
cluster_8
cluster_9
75 28 86 164 13 48 184 136 79 195 35 127 55 152 176 66 133 46 93 61 137 91 177 65 118 11 147 194 179 187 180 89 14 197
-0.8 0 0.8
135 Pou2f2
190 Pou2f2
218 Pou2f2
230 Spi1/Spib/Etv2
205 Bcl11a//Bcl11b
133 Spi1/Spib
255 Spi1/Spib
276 Spi1/Spib
286 Spi1/Spib
122 Ellf1//Nfatc3
120 Ets
11 Ets1
252 Ets
112 Tcf3
8 Zeb1//bHLH
78 Tcf3//Id3
93 Zeb1//Id3//Tcf3
69 Ctcf
275 Ctcf
23 Ctcf
129 Nr2f6
68 Nr1d1//Rorc
220 Nr2f6
238 Nr1d1
257 Pax5
167 Pax5
217 Pax5
97 Eomes//Tbx21
279 Eomes//Tbx21
89 Irf1//Stat2//Prdm1
250 Irf1
94 Spi1
165 Irf1
224 Irf1//Stat2
127 Nfic
40 Nfat
209
34 Cebp
174 Cebp
67 Mzf1
102
194
80 Lef1//Tcf7
166 Lef1//Tcf7
223
173 C2H2 ZF
227
154 Sp
242 Sp//Klf
231 Nfkb-het
247 Nfkb-hom
123 Ebf1
260 Ebf1
b
Filter 174
Filter 183
Filter 34
Second layer filter 48
c
Filter 124
Filter 220
Filter 227
Second layer filter 28
d
Filter 10
Filter 15
Second layer filter 158
e
Filter 288
Filter 106
Second layer filter 38

### Figure S8

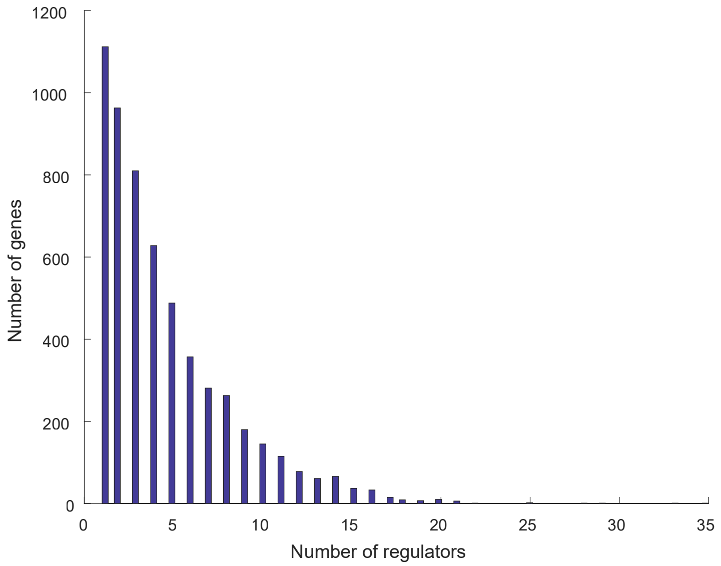

1200
1000
800
600
400
200
0
Number of genes
0
5
10
15
20
25
30
35
Number of regulators
